# A magnetically actuated, optically sensed tensile testing method for mechanical characterisation of soft biological tissues

**DOI:** 10.1101/2022.07.29.501994

**Authors:** Luca Rosalia, Adrien Hallou, Laurence Cochrane, Thierry Savin

## Abstract

Mechanical properties of soft biological tissues play a key role in their normal physiology, contributing to their formation during development, maintenance and repair during adult homeostasis, and driving diseases such as cancer. Mechanics has been proposed to exert its effect by impacting cells fate decisions and cell behaviours including proliferation, differentiation and motility, amongst others. However, despite its critical relevance, a comprehensive analysis of the biomechanics of soft biological tissues is still lacking due to the limitations of the existing characterisation tools. In this article, we describe the development of a device for uniaxial tensile testing of small samples of epithelial and connective tissues, based on the closed-loop interaction between an electromagnetic force actuator and an optical strain sensor. First, we validate the device with synthetic elastomers of known mechanical properties and compare its performance with conventional tensile testing methods; then, we characterise the mechanical properties of the squamous epithelium of the mouse oesophagus along with its supporting connective tissue and underlying muscle in controlled environmental conditions. Through an analysis of strain-stress curves, we demonstrate that the whole oesophagus behaves as a trilayered composite material, whose overall mechanical response depends on the properties of each of its tissue layers. Overall, the proposed setup enables measurements of the mechanical properties of soft biological tissues with unprecedented reliability and precision, and offers an ideal platform for future instrument developments.

## Introduction

Mechanical properties of soft tissues, such as stiffness, strength and viscoelasticity,^1^ are key to numerous biological processes^2^, including embryonic morphogenesis^3–5^, postnatal development^6^, tissue homeostasis^7, 8^, tissue physiological function^9, 10^ and aging^11^. They are also central to the initiation and progression of several pathologies, from cancer^12^, wound healing^13^ and fibrosis^14^, to cardiovascular diseases such as aneurysm or atherosclerosis^15, 16^. Despite this critical relevance, available mechanical data on biological tissues are conspicuously sparse due to limitations in existing characterisation tools and the significant discrepancies that arise between the numerous mechanical testing methods^1, 17, 18^. For example, Young’s modulus measurements of human skin via indentation methods yield values of approximately 35.0 kPa, whereas tensile tests generate values in the 100-200 kPa range^19^. Indentation with a nanoindenter or an atomic force microscope (AFM) is one of the most popular mechanical testing methods for biological tissues, even if it suffers from several limitations. This method involves measuring the penetration depth of a probe into the tissue of interest as a function of the indenter load. The Young’s modulus, or other mechanical properties, are then indirectly evaluated using arbitrary fitting models, specific to the indenter geometry and requiring knowledge of the Poisson’s ratio of the tissue under investigation^20^. Moreover, the method suffers from other significant shortcomings. The load applied to the sample and the resulting elongation are typically in the piconewton and micrometre range respectively. Further, loading is only applied to the superficial layers of the tissue, in a direction perpendicular to the plane of the sample. These mechanical testing conditions are not comparable with those observed *in vivo*, where soft tissues are typically subjected to in-plane millimetric elongations and to mechanical forces in the micro-to milli-Newton range^1^. Uniaxial tensile testing - the gold standard mechanical testing method for engineering materials - provides a way to overcome some of these limitations by allowing direct calculations of the stress-strain response and mechanical properties of the tissue, such as its Young’s modulus, solely by knowledge of the applied load, the elongation of the sample, and its geometry^9, 21^. Moreover, it involves stretching of the sample in a single direction, which can be in the plane of the tissue or perpendicular to it, and in a physiologically relevant range of loads and deformations. Current tensile testing devices for soft tissues are based on customised commercial devices initially designed to test engineering materials^20^ or developed in-house for the particular tissue sample under investigation^16, 21, 22^. Alongside lacking generality, these methods suffer from a variety of limitations, including poor reliability arising from untraceable stress concentrations and assumptions on the geometry and deformation of the specimens, the requirement for large samples prone to exhibiting heterogeneous and anisotropic properties, the inability to replicate the physiological environment of the tissue and to investigate rate-dependence effects.

The lack of appropriate instruments for tensile testing of soft biological tissues, and consequently, of standardised data sets, greatly hinders advances in mechanobiology and their translational applications. To overcome these shortcomings, we describe in this article the development of a tensile testing apparatus that is capable of measuring the mechanical properties of small tissue samples in physiologically relevant conditions. Our device relies on magnetic actuation and optical sensing, enabling simultaneous measurements of the force applied to the tissue, due to the interaction between an electromagnet and a ferromagnetic bead attached to the sample, and of the sample elongation, tracked by an optical sensor. In this work, we first introduce the design and development of our proposed tensile testing apparatus; then, we describe the validation of its performances against conventional tensile testing methods using synthetic elastomers with known mechanical properties. Finally, we provide a direct assessment of the performance of our device, with a study on the mechanical properties of the murine oesophagus and its different constitutive soft tissues, demonstrating the unprecedented accuracy and precision of our method for tensile testing of soft biological tissues.

## Results

### Design considerations for the development of a tensile testing apparatus for soft biological tissues

Prior to proceeding with a detailed description of the design of the proposed opto-magnetic tensile testing apparatus, it is necessary to establish the required operating limits of such a device in terms of the magnitude of forces applied and the range of tissue deformation needed. All of these are primarily determined by the size and mechanical properties of the tissue samples being tested. The main characteristic of our design is its ability to enable testing of soft biological tissue samples at the millimetre scale. This scale corresponds to the typical dimensions of human tissue samples routinely biopsied in clinical settings, as well as to the murine embryonic and adult tissue samples collected for biomedical research. At such scale, the samples are generally smaller than the scale of heterogeneous variations in tissue structure, but large enough to exhibit bulk mechanical properties. Mechanical properties of soft biological tissues in physiological conditions are highly variable, ranging from a Young’s modulus, *E*, of around 0.1 kPa for brain to around 25.0 MPa for ligaments^1^. Considering that biopsied samples range between *d* = 1.0 to 2.0 mm in diameter and 5.0 to 7.0 mm in length, this equates to tissue samples with a longitudinal stiffness 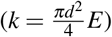 ranging from 80.0 μN for brain tissues to over 20.0 N for ligaments in the linear elasticity regime. Given that most soft tissues have an ultimate tensile strength of around 20.0 MPa, equivalent to force of around 20.0 N, and ultimate tensile strains of around 150 %, equivalent to specimen lengths of 10.0 to 12.0 mm^9^, we conclude that our tensile testing must be able to generate mechanical forces ranging from approximately 10^-6^ to 1.0 N over a range of around 15.0 mm and with micrometre scale spatial resolution.

In a previous study, we demonstrated that a simple magnetic force actuator, composed of a permanent magnet and a steel bead attached to a millimetric sample, can be used to generate forces ranging from 10^-6^ to 10^-3^ N over a range of 2.0 to 8.0 mm from the edge of the magnet, deformation of the sample being tracked using video-microscopy^23^. The force exerted by the magnet on the test specimen is highly non-linear, exponentially increasing as the steel bead approaches the magnet. This unique feature makes magnetic attraction particularly well suited to exert tensile forces on soft tissues, as these tend to exhibit a strain-stiffening behaviour, whereby their stiffness increases non-linearly upon increased deformation^9^. Here, we develop a more versatile, precise and robust tensile testing apparatus, where the attraction between a steel bead attached to the sample and an electromagnet is used to apply a known uniaxial tensile stress, which can be calculated using calibrated force/distance/current characteristics.

In our proposed design, the elongation of the sample is directly tracked by monitoring the shadow of the steel bead on a linear image sensor (CCD). A closed-loop controller modulates the current fed into the magnet, allowing a variety of properties to be measured through both static and dynamic tests, including constant strain, constant stress and oscillatory viscoelastic and fatigue measurements. The advantages of such a design include the large force operating range that can be achieved by altering the current in the electromagnet, position and ball diameter, and the non-contact nature of the magnetic interaction, reducing friction and mechanical complexity. Thus, the architecture of the proposed tensile testing instrument can be split into three sections relating to sample handling, force application and deformation measurement through an opto-magnetic actuator, and test monitoring with image analysis as shown in Figure 1 A.

**Figure 1.**
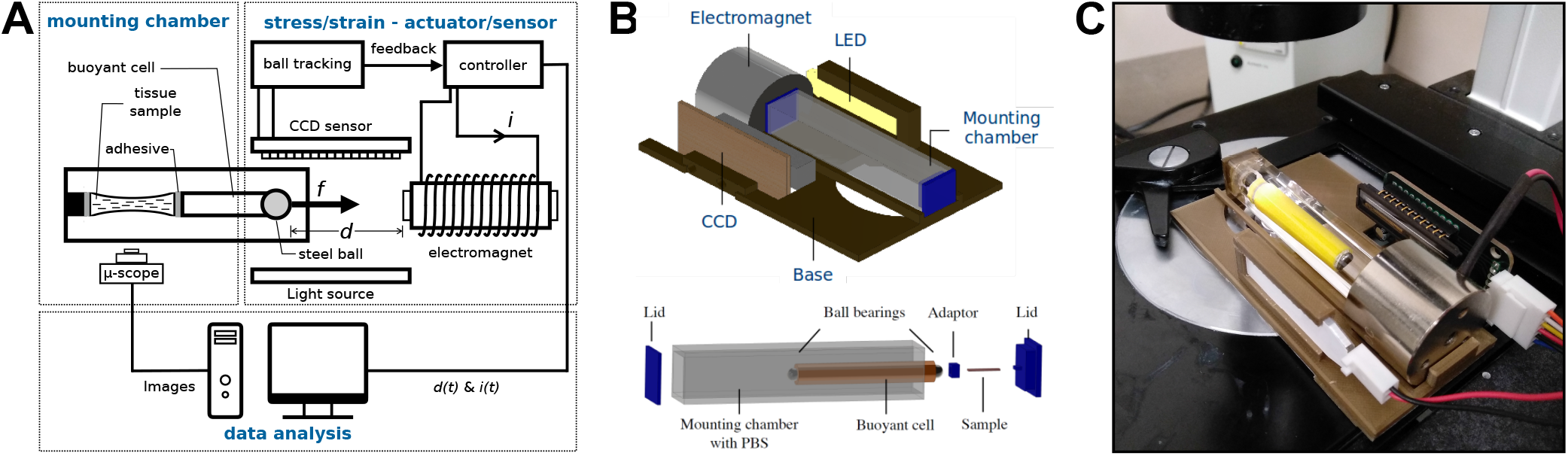
Design of an integrated device for magnetically actuated, optically senses tensile testing of biological tissues. **A**: Schematic illustrating the operation principle of the apparatus designed to allow tensile stress testing experiments of soft biological tissues in the millimetre scale. **B**: 3D rendering of the setup showing (Top) the 3D printed custom microscope stage, the electromagnet, the CCD sensor and the illumination system (LED array); (Bottom) the transparent mounting chamber, buoyancy cell, soft tissue sample, 3D printed lids and sample holders. **C**: Picture of the tensile stress testing apparatus in operation on the stage of an Olympus CKX41 inverted microscope.

### Hardware components and design of a magnetically actuated and optically sensed tensile testing apparatus

The mechanical design of the instrument is centred around the optical setup requirements, aiming to align the light source, CCD, electromagnet and mounting sample chamber in a reliable yet adjustable configuration. The electrical and optical components are integrated with a custom 3D printed base allowing for simultaneous tensile testing and live-imaging of small biological tissue specimens as shown in Figure 1 B and C. 3D-printing was chosen to manufacture the base and various other mechanical part of the device, due to its several advantages over other manufacturing techniques, including low costs, fast prototyping, design customisability and single-step fabrication (cf. Materials and Methods). The electrical hardware is integrated with two printed circuit boards (PCBs). The main PCB, as shown in Figure Sup. 1 A, is designed to attach to the headers of the microprocessor board, allowing interfacing with any of the microcontroller output pins. It contains the electromagnet driver circuit, along with the CCD control circuitry and connections with the current sensor and an external, high-current power source and the electromagnet (cf. Materials and Methods). The CCD is located on a separate “breakout” PCB, which routes the appropriate pins of the chip to a 6-wire parallel bus connection to the main PCB. The use of a separate board aids positioning within the mechanical design, and the stray capacitance between bus wires was found not to adversely affect operation.

The opto-magnetic actuator is composed of two components driven by the microcontroller of the embedded system: a magnetic actuator, consisting in an electromagnet generating a variable magnetic field acting on a ferromagnetic steel bead connected to the tissue sample via a buoyancy cell, and an optical position sensor, composed of a uniform light source casting the shadow of the steel bead on the CCD with a resolution of *δx* = 106.0μ*m*, a precision of 1.6*μm* and a tracking speed of 200 Hz (cf. Materials and Methods). The pull force *F_E_* generated by the electromagnet on the steel bead is dependent upon the diameter *d* of the ball, on its distance from the magnet x, and on the current supplied *I*, as per the phenomenological relation 1:

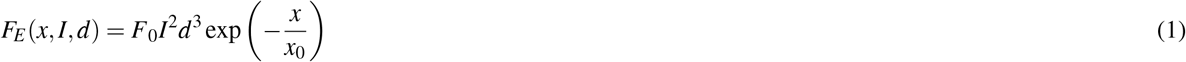

where *x* is the distance from the surface of the magnet to the centre of the ball, and 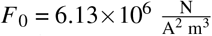 and *x*_0_ = 2.258 ×10^-3^m are constants (cf. Materials and Methods and Figure Sup. 1 E and F). Calculation of the attractive magnetic force is therefore enabled by real-time measurements of the electromagnet current and of the distance between the electromagnet and the bead by means of the CCD. These are interfaced with the microcontroller of the embedded system, which offers serial-over-USB functionality for communication with an external PC, enabling both definition of the test parameters and data collection during the experiments.

### A mounting chamber and buoyancy cell allow for tensile testing in physiological conditions

One of the most critical requirements for mechanical testing of soft biological tissues is the need to maintain them as closely to their physiological conditions (temperature, pH, osmolarity, etc.). To this aim, we designed a mounting chamber that allows the immersion of the tissue test specimen in a phosphate-buffered saline (PBS) solution or culture medium. To minimise any interference with the optical system, the mounting chamber was chosen to be transparent and of square cross-section. A neutrally-buoyant cell was devised to reduce gravity-induced noise and guarantee the alignment of the steel bead with the CCD sensor. The cell was designed to be hollow, its dimensions being computed to match the density of PBS, and manufactured in PLA via 3D-printing. A cylindrical shape was employed for the cell to curtail drag effects and provide symmetry with respect to three mutually-perpendicular axes, preventing the system from rotating. For this purpose, one ball bearing was fixed at each end of the cell, the closest of which being attracted to the electromagnet. From Equation 1, the interaction between the other bead and the magnetic field was computed to be several orders of magnitude smaller, due to its greater distance from the magnet, and was thus considered negligible. Instead, this ball bearing was used as an attachment site to the sample. The mounting chamber and the neutrally-buoyant cell, together with the lids, the adaptor, and an illustration of the specimen, are depicted in Figure 1 B and C.

Analysis of the drag effects on the buoyancy cells revealed close agreement between the experimental results and the numerical simulation (cf. Materials and Methods), as depicted in Figure Sup. 4 in Supplementary Information. In accordance with the fluid dynamics theory of simpler shapes^24^, we found that the drag force increased linearly with velocity at small Reynolds numbers, and that smaller cross-sectional areas of the chamber yielded a higher flow resistance. The latter effect arose from an increase in the blockage ratio *β*, i.e. that is between the diameter of the object and the side length of the channel^25^. An expression of the drag force F_D_ as a function of velocity *u* was therefore obtained upon fitting of the experimental data to a straight line via linear regression using the least-square method (R^2^ =0.9789). We report here, the result obtained for 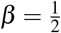, which corresponds to the design we have used for all the testing experiments described in this article:

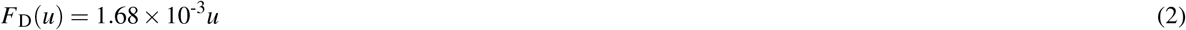

The intercept was in the order of *μ*N, and it was thus considered negligible. By means of this empirical model, we concluded that for a broad range of physiological strain rates 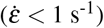, the drag force, *F*_D_ ≪ 0.050 *μ*N, is negligible in comparison with the force required for stretching biological specimens at the millimetre scale, even for the softer tissue samples, as previously outlined.

### A closed-loop feedback controller enables reliable characterisation of soft tissue mechanical behaviours

An analysis of a mechanical model of the device and of its open-loop behaviour highlights the inherent instability of the interaction between the ferromagnetic bead and the electromagnet, which can lead to the ball accelerating exponentially towards the magnet, a phenomenon herein referred to as “pull-in” and happening for deformations as small as 30 % (cf. Supplementary Information and Figure Sup. 5 A-D). It was thus necessary to integrate in our design a closed-loop feedback mechanism to ensure that stability is achieved under the effects of unknown sample mechanical properties (these are the mesurands), as well as disturbances such as external vibrations, sensor noise and sensor delays.

We implemented an adaptive proportional-integral-derivative (PID) embedded controller whose block diagram is shown in Figure Sup. 5 E and whose mathematical model is derived in the Supplementary Information. To define numerical values for the closed-loop poles of the system, and thus the gains of the adaptive PID controller, we simulated the behaviour of the system via a *Simulink* model depicted in Figure Sup. 5 F. The closed-loop poles were selected to be p_1_ = −60, p_2,3_ = −6.5 ± j (cf. Supplementary Information). This first pole value allows the integrator to quickly adapt to the unknown sample force, while the complex conjugate pair of poles gives a reasonably fast but well damped and low overshoot transient response. To maintain stability at high sample deformations in spite of the delay in CCD measurements, we implemented an adaptive controller, whereby the gains are dynamically recalculated to give unchanging closed loop poles as, the magnetic spring constants, *k_x_* and *k_I_*, vary with the bead position *x* (cf. Supplementary Information).

Last, to prevent integrator wind-up for samples displaying high damping constants *b*, we implemented the following bound on the integrator state, 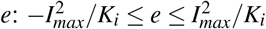, where *I_max_* is the maximum current value and *K_i_* is the integral gain of the controller (cf. Supplementary Information). Overall, as depicted in Figure Sup. 5 G, despite a slight increase in the amplitude of the oscillation observed at the beginning of the test, the selected poles allow an excellent controllability of the system, with errors never exceeding 10%. This was further confirmed by our experimental validation as shown in Figure Sup. 5 H.

### Testing of synthetic materials with known mechanical properties validates device performance

Validation of our device allowed us to confirm that it is suitable for mechanical testing of soft biological tissues and to assess its performance against established instruments. As shown in Figure 2 A, Polyvinyl siloxane (PVS) specimens, made of Elite Double 8 and 22 (cf. Materials and Methods), were tested on our proposed device and on an Instron tensile testing machine, which represents a gold standard tool for mechanical characterisation of engineering materials. PVS, which is a widely used silicone elastomer was chosen for its relatively low stiffness range, that is comparable to that of numerous soft biological tissues. Analysis of the linear elastic behaviour of these samples (cf. Materials and Methods) demonstrated adequate functioning of the instrument, as no major differences could be observed upon comparison with a well-established method for uniaxial tensile testing. Figures 2 B and C illustrate the mechanical response to 10% strain of the Elite Double 8 and 22 specimens, as tested on our proposed instrument, either in air or in the mounting chamber filled with PBS, and on an Instron machine. Measurements of the Young’s modulus, *E*, were obtained upon computation of the slope of the stress-strain curves at 5-10% elongations and are summarised in Table 1. These data demonstrate that the mechanical responses recorded by our device are highly accurate and in close agreement not with those measured by the standard method, as well as with the values of *E* provided by the manufacturer of this synthetic elastomer.

**Figure 2.**
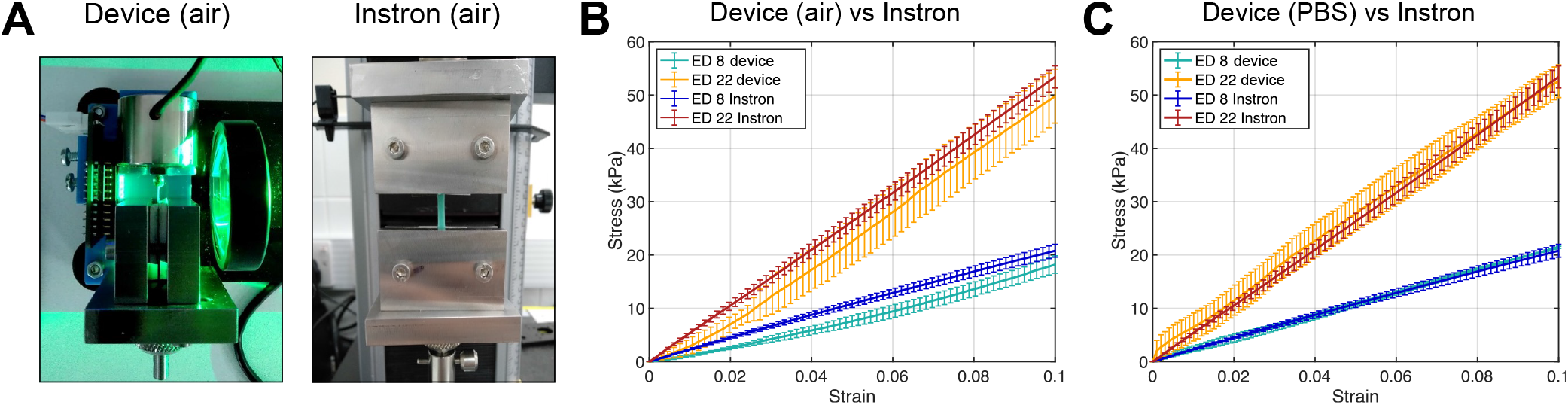
Device validation on synthetic material specimens: mechanical characterisation of PVS samples and comparison with an Instron machine. **A**: (Left) Picture of the tensile stress testing apparatus in operation (vertical configuration) without the mounting chamber and using a metallic clamp to mount an elastomer test specimen. (Right) Picture of an Instron machine with an elastomer test specimen mounted with metallic clamps. **B**: Stress-strain curves of Elite Double 8 and 22 samples on our device (vertical configuration - air) and on an Instron machine respectively. N=3 samples per condition. **C**: Stress-strain curves of Elite Double 8 and 22 samples on our device (horizontal configuration - mounting chamber filled with PBS) and on an Instron machine respectively. N=3 samples per condition. Error bars represent 1 SD.

**Table 1.** Young’s modulus of the Elite Double 8 and 22 PVS specimens measured on our device, on an Instron tensile testing machine, and as provided by the manufacturer. Experimental value are shown as mean ±1 SD.

| Method | $E$ (kPa) | |
| --- | --- | --- |
|  | Elite Double 8 | Elite Double 22 |
| Device (mounting chamber - PBS) | $216.0 \pm 11.0$ | $541.8 \pm 18.4$ |
| Device (no mounting chamber - Air) | $213.8 \pm 15.6$ | $545.7 \pm 35.8$ |
| Instron 5544 | $198.9 \pm 10.9$ | $542.8 \pm 10.5$ |
| Literature | 211 | 562 |

When clamps were used to mount the test specimen (Figure 2 A, air configuration), the stress-strain response exhibited a slightly non-linear behaviour at low strains. This was particularly apparent on the Elite Double 8 test specimens, as shown in Figure 2 B. Using finite element modeling (cf. Materials and Methods), we demonstrated that such non-linearity can arise from the torsion of the test specimen around the tensile axis. As evidenced in Figure Sup. 6 A and B, even relatively small twist angles can generate non-linear stress-strain responses. Using video-microscopy to capture the specimen dynamics during testing, we discovered that torsion was generated by an asymmetric attachment of the sample on the steel bead resulting in an oscillating motion of the sample due to gravity. This hypothesis was corroborated by conducting tensile testing experiments with the device switched in the vertical direction so that gravity acted uniaxially to the electromagnetic force, preventing oscillations in the normal direction, and by designing a 3D printed mounting adapter and mounting support (Figure Sup. 6 D) allowing an easy and symmetric attachment of the test specimen. Results depicted in Figure Sup. 6 C, demonstrate the validity of both the hypotheses, as the measurements obtained on the vertical setup were affected by only minimal noise, while use of specimen-to-ball adaptors completely eliminated the non-linear bias at small strains.

To overcome this limitation, and taking into account the need to maintain biological samples in a mounting chamber filled up with a physiological medium, we incorporated in our design the cylindrical neutrally-buoyant cell described in the previous section and shown in Figure 1 B and C. Using the same testing protocol, each PVS sample being tested three times up to *ε* =0.1 at a constant strain rate 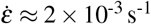, we observed that our device yielded measurements highly consistent with those given by an Instron machine as shown in Figure 2 C and reported in Table 1. To further assess the robustness of the measurements generated by our apparatus, we evaluated hysteresis (i.e. the dependence of output values on the system history) by monitoring the stress-strain behaviour of Elite Double 8 sample in loading and unloading states. The specimen was stretched and relaxed at the same rate 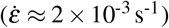 following a triangular input signal. Three repetitions yielded the results in Figure Sup. 6 E, which show negligible difference between the upper (loading) and lower (unloading) curves, demonstrating that the device is not affected by hysteresis.

### Tensile apparatus allows biomechanical testing of the murine oesophagus over a range of testing parameters

To validate the performance of our proposed tensile testing apparatus on soft biological tissues, we next studied the mechanical properties of the oesophagus. Propelling food boli from the pharynx to the stomach, the oesophagus is an organ with a profound biomechanical function^9, 10^, and recent works have unravelled that knowledge of its mechanical behaviour from the organ^26^ to the cellular level is paramount to comprehend its development,homeostasis, physiology and remodelling during disease^6, 27, 28^. Furthermore, its constitutive tissues are representative of the soft tissues encountered in most mammalian organs: epithelia, extracellular matrix (ECM) rich connective tissues and muscles. A tubular-shape dorsal organ from the upper gastrointestinal tract whose anatomical structure and location are similar in mouse and human, the oesophagus is composed, as shown in Figure 3 A, of multiple tissue layers surrounding a hollow central lumen: the mucosa, submucosa and tunica muscularis^29^. The mucosa is constituted of a squamous stratified epithelium with several layers of differentiated suprabasal cells and one layer of self-renewing basal progenitor cells^6^. The submucosal or stroma contains the lamina propria - which underlies the epithelium and consists mainly of ECM rich connective tissue - and the muscularis mucosae. The externally-apposed tunica muscularis is constituted of the circular inner and the longitudinal outer muscle layers. To date, a comprehensive characterisation of the mechanical behaviour of the oesophagus is missing due to the absence of adequate testing methods and protocols for tissue separation of multi-layered biological organs. Past experimental analysis has been confined to opening-angle experiments or inflation tests to measure circumferential strains in a variety of non-model systems: rat^30–32^, rabbit^33, 34^, guinea pig^35^ and pig^36, 36–38^, with no available data from tensile testing studies in mice or human.

**Figure 3.**
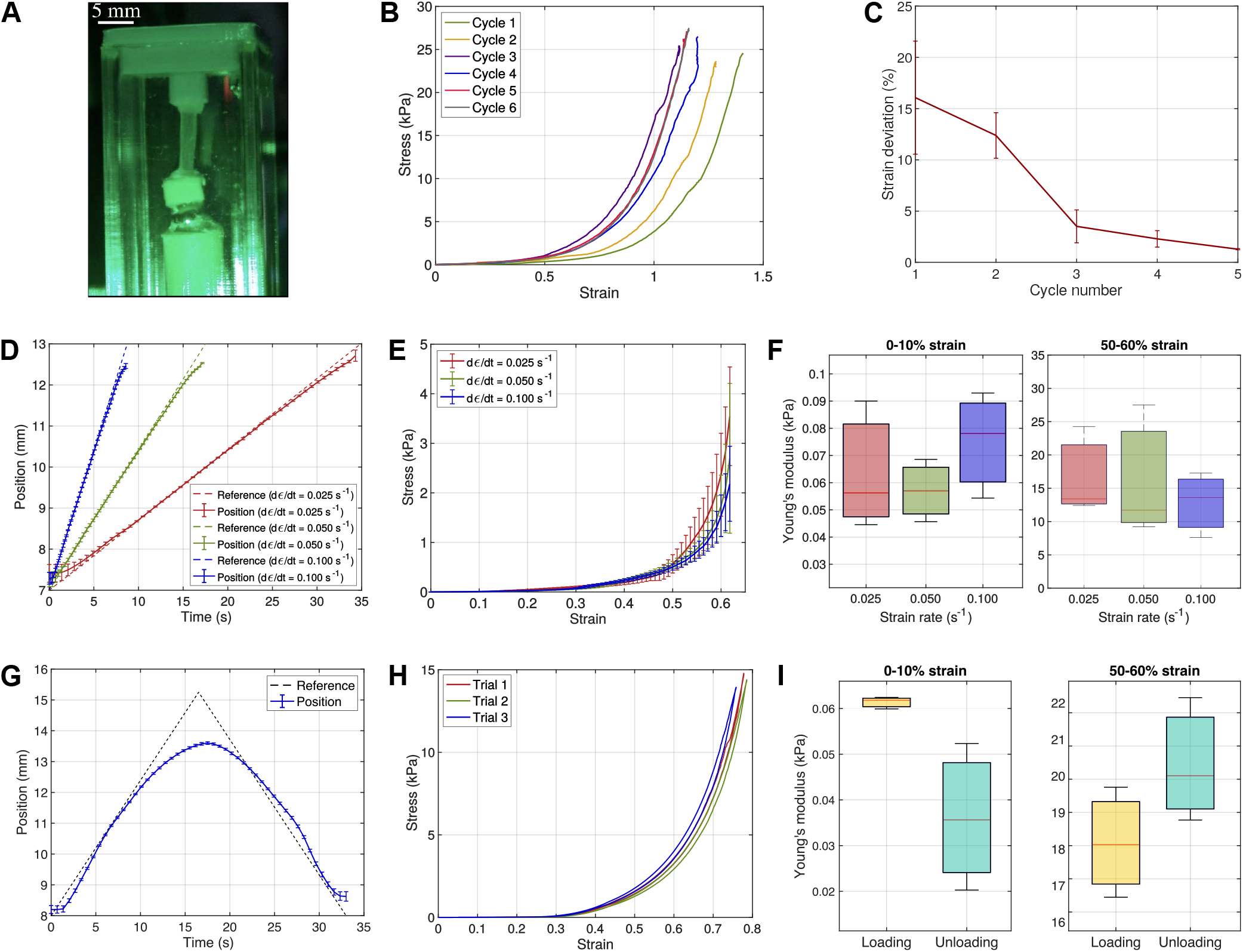
Tensile testing of the murine oesophagus over a broad spectrum of testing parameters. **A**: Lateral view of an oesophagus wall test specimen under uniaxial tension during testing on our proposed device. **B**: Stress-strain behaviour of an oesophagus test specimen for preconditioning analysis over 6 consecutive loading cycles. **C**: Absolute strain deviation from the mean value of the subsequent loading cycles for each repetition in respect of the number of loading cycles. **D**: Reference input tracking for strain rate analysis of oesophagus wall test specimens as performed for 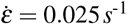,0.050s^-1^, and 0.100s^-1^. **E**: Stress-strain behaviour of oesophagus samples for different strain rates. **F**: Young’s modulus at 0-10% and at 50-60% elongations for strain rate analysis. **G**: Reference input tracking for hysteresis analysis for oesophagus test specimens. **H**: Stress-strain behaviour of oesophagus samples during loading and unloading cycles for hysteresis analysis. Error bars represent 1 SD. **I**: Young’s modulus at 0-10% and 50-60% elongations during hysteresis cycle. Error bars represent 1 SD.

Prior to investigating the mechanical behaviour of the oesophagus and its constitutive tissue layers, the effects of preconditioning were examined due to the absence of a standardised protocol. Preconditioning consists in repeating load cycles to obtain a steady and repeatable mechanical response of the test specimen and is a common procedure in testing of soft biological tissues^21^. We observed the variability associated with the mechanical response of one sample for each individual layer over multiple preconditioning cycles. Figure 3 B illustrates the mechanical response of the epithelial layer to multiple loading-unloading cycles, while Figure 3 C shows the absolute strain deviations averaged across the three tissue layers at each cycle. This was calculated at the point of maximum common stress and with respect to the mean value of the subsequent repetitions. Results demonstrated that three preconditioning cycles were sufficient to reduce the strain deviation below 5%.

The stress-strain behaviour of single specimens of oesophageal wall was measured at different strain rates. This was paramount in that rate sensitivity is known to vary considerably among different tissues^9^ and allows to determine the optimal rate value for quasi-static mechanical testing to probe the elastic behaviour of the tissue. The test followed three preconditioning cycles and consisted of three repetitions for each strain rate being employed (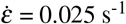, 0.05 s^-1^, 0.1 s^-1^). Although in the majority of experimental studies this is based on standard protocols for engineering materials and corresponds to values of approximately 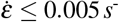^16^, our analysis revealed that no significant difference arises at 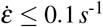. Results are depicted in Figure 3 D, E and F which shows the excellent controllability achieved over the given range of strain rates, the mechanical response of the oesophagus wall, alongside the Young’s modulus calculated at both low (0-10%) and high (50-60%) deformations.

Differences in the response of the oesophageal tissue during loading-unloading cycles were subsequently investigated by conducting hysteresis analysis on individual test specimens. For this purpose, a triangular input was used as the reference position signal as illustrated in Figure 3 A. Tests were carried out at 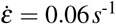 and repeated three times on each single specimen. Inspection of the mechanical response (Figure 3 H) and measurements of the Young’s modulus at low and high strains (Figure 3 I), showed only marginal variation between the loading and unloading phases. During unloading, the measured stress decreased more rapidly than in the loading phase. As a consequence, the tissue was slightly stiffer at high strains and more compliant at low strains during the second phase of the hysteresis cycle. Overall, strain-rate and hysteresis analyses demonstrated that time-dependent effects were negligible for preconditioned tissue samples, further validating the capability of our device to be able to isolate the elastic behaviour of the soft biological tissues under investigation.

### Device performance allows the characterisation of the oesophagus as a multi-layered soft tissue organ

Based on findings from our previous analysis, preconditioning was conducted and a strain rate of 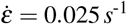 was prescribed. Figures 4 B and C illustrate the behaviour of the intact oesophageal wall, alongside that of each separated tissue layer in both the 2- and 3-layer configurations respectively. These plots highlight the reproducibility of the results generated by our proposed device, with errors in the stress-strain response generally falling below 15%. This constitutes one of the major advantages of the device and method presented in this article and is unprecedented in the literature. Although the inter-sample variability was negligible for any of the tissues under investigation, greater divergences arose in the inter-sample deviation across the specimens, with slightly larger error being associated with the separated tissue layers than with the intact oesophagus. This could be explained by virtue of the methods used to separate the different tissue layers, which may be responsible for inducing damage or undesirable pre-stress within their structure.

**Figure 4.**
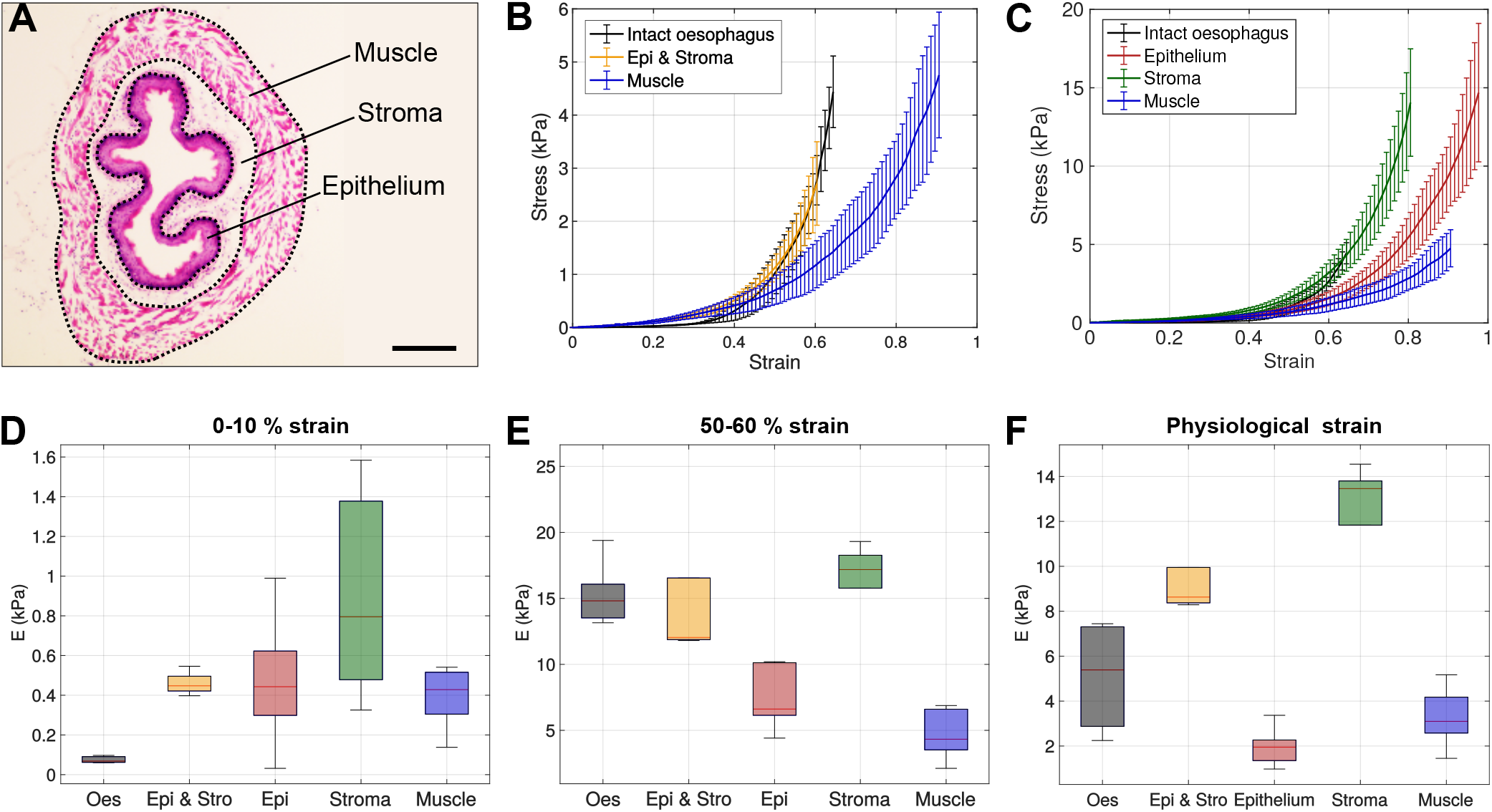
Mechanical characterisation of the oesophagus as a multi-layered soft tissue organ. **A**: H&E stained section of the oesophagus showing the epithelial (mucosa), stromal (submucosa) and muscle (tunica muscularis) layers. Scale bar is 200 μm. **B**: Stress-strain behaviour of the intact oesophagus, undivided epithelium and stroma, and muscle layer. Error bars represent 1 SD. **C**: Stress-strain behaviour of the intact oesophagus, and of the epithelial, stromal, and muscle layers. Error bars represent 1 SD. **D**: Young’s modulus at 0-10% elongations of the intact oesophageal wall and of the separated tissue layers. **E**: Young’s modulus at 50-60% elongations of the intact oesophagus and of the separated tissue layers. **F**: Young’s modulus of the intact oesophagus and of the separated tissue layers at physiological strain.

In the 2-layer configuration, despite being more compliant at low strains, the inner layer (epithelium and stroma) was significantly stiffer than the outer layer (muscle) at elongations superior to approximately ε =0.45. This result is close to the measured in vivo longitudinal strain ε_S_ =0.43 ±0.06 and comparable with that reported for inflation tests, where crossing occurs at radial strains close to ε = 0.5^30^. In the 3-layer configuration under the same stress-free conditions, it is apparent that the stromal layer exhibits the stiffest behaviour, dominating the response of the entire oesophagus wall. This likely arises from the increasing stiffness exhibited by the ECM rich stromal tissue following uncrimping of collagen fibres^39^. Contrarily, the muscle layer was the most compliant layer, with the response of the epithelium being intermediate between that of the muscle and of the stroma. This was confirmed by computing the Young’s modulus, as shown in Figure 4 D and E. At high strains (50-60%), the Young’s modulus of the stroma dominates by a factor 2-3 those of the epithelium and the muscle. However, at low strains (0-10%), the intact oesophageal wall exhibited the lowest stiffness, while the different tissue layers were displaying relatively similar Young’s modulus values, all below 1.0 kPa.

Albeit informative of the response of each tissue layer, values of the Young’s modulus of the oesophagus wall in the intact state cannot be compared with measurements obtained for each layer in the zero-stress state at equal elongations. Indeed, this does not represent a reliable indication of the relative contribution that each tissue layer provides to the oesophagus wall under physiological conditions, due to the presence of non-zero residual strains between tissue layers. As per Equation 30 in Supplementary Information, evaluation of the contribution provided by each tissue layer in the physiological state requires the knowledge of their thickness fractions *τ* and residual strains ε_R_ (cf. Supplementary Information). We thus measured the thickness distributions of the different tissue layers on oesophagus tissue sections, as well as the residual strains in the longitudinal direction (cf. Materials and Methods), as summarised in Table 2. Then, we computed the Young’s modulus of the oesophagus wall and each tissue layer at physiological strain *ε_P_* = *ε_R_* + *ε_S_* (cf. Supplementary Information) and reported the results in Table 2. As illustrated in Figure 4 F, this analysis suggests that, at physiological strain, the response of the intact wall is dominated by the stromal tissue layer. As summarised in Table 2, these results are validated by our model (cf. Supplementary Information), which closely predicts the behaviour of the intact oesophageal wall for both the 2- and the 3-layer configurations.

**Table 2.** Thickness, *h*, longitudinal residual strain, ε_*R*_, *in vivo* longitudinal strain, ε_*S*_, longitudinal physiological strain ε_*P*_ and Young’s Modulus at physiological strain, *E_P_*, of the oesophagus and the different oesophageal tissue layers. Strains are computed using the cut open oesophagus wall as the reference configuration. For the oesophagus, the direct experimental measurement is compared with the semi-analytical predictions of the 2- and 3-layer models. Values are shown as mean ±1 SD.

| Tissue | $h$ ($\mu m$) | $\epsilon_R$ | $\epsilon_S$ | $\epsilon_P$ | $E_P$ (kPa) |
| --- | --- | --- | --- | --- | --- |
| Oesophagus (Experimental) | $312.44 \pm 11.44$ | 0.00 | $0.43 \pm 0.06$ | $0.43 \pm 0.06$ | $5.11 \pm 2.32$ |
| Oesophagus (2-layer model) | — | — | — | — | $5.06 \pm 1.04$ |
| Oesophagus (3-layer model) | — | — | — | — | $4.22 \pm 0.74$ |
| Epithelium & Stroma | $83.44 \pm 5.57$ | $0.06 \pm 0.04$ | — | $0.49 \pm 0.07$ | $9.68 \pm 2.31$ |
| Epithelium | $45.6 \pm 5.53$ | $-0.21 \pm 0.08$ | — | $0.22 \pm 0.10$ | $1.98 \pm 0.82$ |
| Stroma | $37.84 \pm 0.68$ | $0.02 \pm 0.12$ | — | $0.45 \pm 0.13$ | $12.22 \pm 3.05$ |
| Muscle | $229.03 \pm 9.96$ | $-0.02 \pm 0.02$ | — | $0.41 \pm 0.06$ | $3.26 \pm 1.29$ |

Overall, testing of the oesophageal wall and its different constitutive tissue layers allowed us to validate the ability of our proposed tensile testing apparatus to provide reliable and highly accurate measurements of the elastic properties of the most common biological soft tissues.

## Discussion

In this article, we discuss the development and characterisation of a device for uniaxial tensile testing of soft biological tissues. The proposed actuation and sensing system is based on the interaction between an electromagnet and a ferromagnetic bead on one hand, and on the coupling of a light source with an image sensor for position tracking on the other hand. Our device was first demonstrated to be suitable for measurements of the elastic properties of synthetic materials and shown to be in close agreement with those measured by means of current gold standard methods in the field. A mounting chamber was then integrated within the device to allow testing of biological specimens in a controlled environment to mimic their *in vivo* conditions as closely as possible and preserve their biomechanical properties. This was coupled with a neutrally-buoyant cell to minimise the effects of gravity, which could otherwise constitute a significant source of measurement noise. A protocol for sample preparation and installation onto the device was outlined to ensure correct alignment, eliminating any torsion and shear effects. Following experimental and numerical analysis of the drag force acting on the neutrally-buoyant system during testing, a generalised mechanical model of the device was formulated and an embedded control system based on an adaptive gain PID controller was implemented to enhance performance and testing range of the device.

The proposed apparatus was then leveraged to investigate the biomechanics of a whole organ, the murine oesophagus, and its different constitutive soft tissues. For this purpose, the optimal number of preconditioning cycles prior to testing was evaluated, and optimal values of the strain rate were determined for isolation of the elastic behaviour of the tissues under investigation. Further analysis demonstrated that, in quasi-static conditions, only minor differences arise in the response of oesophageal tissues between the loading and unloading phases of its hysteresis cycle. Upon separation of each tissue layer, we conducted the first biomechanical analysis of the oesophagus as a multi-layered structure. While the stroma exhibited the stiffest response at high and physiological strains, the muscle layer was shown to provide the major mechanical contribution to the oesophageal wall at low-to-intermediate elongations in the stress-free state. Contrarily,the epithelium exhibited an intermediate behaviour in the former state, while it was found to be the most compliant layer of the undivided wall at physiological strain.

This investigation corroborated the high reliability of the proposed device, which yielded errors in the stress-strain response below 15%. This represented a considerable improvement from a former device developed by this group based on an open-loop permanent magnet, whose measurements were affected by inaccuracies ranging from 20 to 30%^23^, and establishes a new standard of precision in the field. Furthermore, such a level of accuracy was key to be able to distinguish the difference of mechanical behaviour of the different tissue layers of the oesophagus in different regimes of force and deformation, and could be used to resolve the heterogeneity of the mechanical properties of other tissues and organs.Overall, our proposed instrument was shown to successfully address the main limitations which other mechanical testing devices suffer from, providing measurements with greater reliability and precision. Moreover, as it allows to test any soft tissue in conditions close to their physiological environment, it may constitute a new standard in the field and a unifying tool for biomechanical characterisation of biological tissues.

Future investigations will aim to further leverage the controllability of our device in a large range of forces and strains to focus on the characterisation of the viscoelastic behaviour exhibited by soft biological tissues via creep and stress relaxation experiments,as well as dynamic testing for evaluation of tissue fatigue. Second, taking advantage of our design, which potentially allows integration with a confocal or multiphoton microscope, we endeavour to couple mechanical characterisation of soft tissues with live imaging of their ECM organisation using Second-Harmonic Generation (SHG)^40^ and of their cellular organisation and behaviour using genetically encoded fluorescent reporters^41^. Further, coupling our device with light-sheet and/or superresolution microscopy could allow us to probe, at the molecular level, the real-time response of the nucleus, organelles, cystoskeletal filaments or even individual transmembrane proteins, ultimately elucidating mechanisms by which cells in a tissue are able to sense and respond to mechanical stimuli. The present design could be modified to tightly control the temperature, CO_2_/O_2_ and chemical composition of the medium in the mounting chamber, paving the way towards more physiologically relevant investigations of mechanical properties and biological behaviours such as gene expression, cell division, cell fate decisions, tissue growth or ECM remodelling, in response to both biochemical and mechanical perturbations. Finally, in future studies, we aim to investigate the mechanical behaviour of a broader range of mouse and human soft tissues, leveraging our device to generate the first “mechanome” - a universal and comprehensive characterisation of soft tissues mechanical properties in homeostasis and disease. Through analysis of the biomechanics of healthy tissues and their changes as they occur during disease, our device could eventually identify alterations in tissue properties of diagnostic or prognostic relevance. This could ultimately support physicians in making informed and individualised treatment decisions and contribute to the translation of mechanobiology research into the clinical arena.

## Materials and Methods

### 3D printed hardware

The base of the device and the mechanical parts such as the mounting chamber lids, the tissue gluing adapters or the tissue mounting support were 3D printed in Polylactic acid (PLA) due to the good mechanical and thermal properties of this material. An Ultimaker 2+ printer (Ultimaker) was used for 3D printing with a layer resolution of 0.1 mm under standard operating conditions. All the blueprints, CAD files and printing files of the different 3D printed parts can be found in Supplementary Information.

### Mounting chamber and buoyancy cell

The mounting chamber was manufactured via laser-cutting of a clear thin-walled hollow polymethyl methacrylate (PMMA) tube (Alternative Plastics). PMMA was chosen for its low cost, ease of processing and biocompatibility. The length of the chamber (76 mm) was determined to accommodate the neutrally-buoyant cell and allow for the desired elongations of the specimen during tensile testing. Custom lids were 3D printed in PLA to provide sealing at each end. The buoyancy cell was designed to be a hollow cylinder, its dimensions and the thickness of its wall being computed to match the density of PBS (cf. Supplementary Information). Similarly to other mechanical parts of the device, the buoyancy cell was manufactured in PLA via 3D printing.

The drag force *F*D acting on the buoyancy cell in the mounting chamber was evaluated both experimentally and numerically. To experimentally measure the drag force, the weight of the neutrally-buoyant cell was reduced by thinning of the inner wall. The cell was first brought to the bottom of a vertically-rotated mounting chamber by means of a permanent magnet and then released to float. Analysis was repeated for two sizes of the mounting chamber to investigate the effect of different cross-sectional areas. The drag force was calculated upon knowledge of the inertia and of the volume of the body, and position measurements obtained by means of a high-speed camera and of Tracker 5.0 motion analysis software.

FEM was conducted on Abaqus to validate the experimental results via fluid-structure interaction (FSI) co-analysis of the modelled cell and the medium. For this purpose, non-zero velocities were defined for the neutrally-buoyant object, while no-slip boundary conditions were imposed at the walls of the medium. Hence, the resultant forces in the direction opposite to motion were computed.

### Magnetic force actuator

A commercial electromagnet (Eclipse Magnetics M52173) was used as the actuator of the apparatus. The number of turns of wire, N, in the coil was determined by matching the simulated inductance to that measured experimentally. The inductance was measured to be 63.2 mH allowing N to be calculated as 675 turns. The resistance of 39.0 **Ω** was matched by adjusting the thickness of the wire to 36 SWG. The rated operating voltage and current are 12.0 V and 280.0 mA, respectively. A FEM of the magnetic field generated by the coil, as shown in Figure Sup. 1 B, was created using FEM Magnetics 4.2 software. Using this model, it was possible to evaluate the magnetic force acting on 4.0 mm diameter grade 1000 hardened carbon steel bead (Simply) by numerical integration of the Maxwell Stress Tensor over its volume^42^, and to generate a set of characteristic curves representing either the force acting on the bead as the function of the current or the force acting on the bead as a function of the distance to the electromagnet. FEM predictions were validated with experimental measurements of the magnetic force acting on the steel bead using a microbalance with a resolution of 1.0 mg, equating around 10^−5^ N, while the position was adjusted using a micrometre head, with a vernier caliper scale resolution of 10.0 μm (cf. Figure Sup. 1 C and D).

### Optical sensor

Measurement of the elongation of the sample was carried out using the shadow cast by the steel bead and buoyancy chamber on the CCD. The CCD (Toshiba TCD1304DG) consists of 3648 pixels at 8.0 μm spacing. Creating a collimated beam of light was necessary to cast sharp shadows and to avoid parallax errors due to the distance between the ball and the CCD. To do so, we used two approaches, the first involving use of a LED panel (RS Components) as shown in Figure 1 B and C, allowing for a compact design of the device, and a second approach which leveraged a combination of a point light source and a positive focal length lens (Figure Sup. 3 A) to further enhance the resolution. Using the CCD output, as shown in Figure Sup. 3 B, the resolution of the sensor, was computed to be *δx*=115.0 *μ*m or 14 pixels in the first case and *δx*= 106.0 *μ*m or 13 pixels in the second. This demonstrates that the resolution of the sensor is optics-limited and validates the use of the LED panel design to improve the compactness of the device and integration on a microscope stage.

As shown in Figure Sup. 3 C, the driving and readout of the CCD array are directly implemented using the MCU timers, analogue to digital converters (ADC) and direct memory access bus (DMA). The master clock is implemented using a 16 bit timer, used in pulse width modulation mode with a duty cycle of 50% and set to a period of 21 core clock pulses to generate a 4 MHz square wave from the 84 MHz reference. The integration clear gate (ICG) is implemented with a 16 bit timer in PWM mode and a pre-scaler of 42 to generate a 2 MHz intermediate base reference. The period is set to 40, equating to a 20 μs CCD integration time. For the ICG, a 32 bit timer is used due to the relatively long period between frame capture events, resulting in a frame rate of 200 Hz. Measurement of the analogue output stream of the CCD requires synchronous operation of the ADC and automatic handling of the sampled values. This is achieved by a system of four interconnected hardware interrupts, an ADC Trigger and the DMA. The ADC Trigger timer is set to 1 MHz, i.e., the CCD data output rate, and generates interrupts that initiate an ADC conversion with eight bit precision. Upon completion of the conversion, an interrupt generated by the ADC triggers the DMA to transfer the sampled value to a buffer array, the same length as the CCD signal. Once the buffer is filled, a DMA Transfer Complete interrupt stops the ADC trigger timer, preventing further samples from being converted, and initiates the execution position measurement algorithm and the next time step of the feedback controller. This cycle is re-initiated by an interrupt generated at the end of an ICG pulse, which re-starts the ADC trigger timer at the same time as the next sequence of CCD output data begins thus continually providing the latest CCD data in the buffer array.

### Controller

A derivation of a generalised model of the system and an analysis of its open-loop behaviour are given in Supplementary Information. An adaptive proportional-integral-derivative (PID) embedded controller was implemented and the software *Simulink* was used to simulate its behaviour in response to a variety of reference input signals to determine its close-loop poles and the gains of the different controllers. The *Simulink* model is composed of 6 main components and described in detail in Supplementary Information. Briefly, the electromagnet block is based on the expression for *F_E_* given in Eq.1, along with the transfer function from duty cycle to current of the MOSFET-based switching circuit which drives the electromagnet. The sample block matches a Kelvin-Voigt model with a bias force, and the inertia of the ball bearing is described by a double integrator, with the output being limited to the dimension of the mounting chamber. The drag block is based on the expression for FD given in Eq.2 and allows computation of the net force acting on the sample. Accurate models of the current and CCD sensors have also been used, including the effect of noise, delay and quantisation. An arbitrary microcontroller was implemented by a MATLAB function block with inputs of the measured current, measured position, and a reference signal, and returning the PWM duty cycle as the output. Stiffness values of 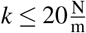 were used in the simulations to mimic the behaviour of stiff biological tissues at 100% strain. Experimental validation of numerical results was obtained by testing Elite Double 8 specimens using a ramp as the reference input position signal to be tracked 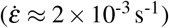. Three repetitions of this experiment were conducted for averaging.

### Embedded system

The embedded design of the device is centred around a ST Microelectronics NUCLEO-F103RB board. Its 84 MHz ARM M4 core is used to implement the feedback controller, while its microcontroller unit (MCU) interfaces with the electromagnetic force actuator drive circuit and the CCD based optical sensor. Embedded programming is undertaken in C using the Hardware Abstraction Layer firmware provided by ST Microelectronics. Design of the feedback controller and optical sensor embedded systems are detailed in Supplementary Information. External data acquisition is enabled by a PC virtual terminal using the software PuTTY and a serial link over USB functionality (virtual COM port).

### Synthetic material samples preparation, testing experiments and simulations

Elite Double 8 and Elite Double 22 solutions (Zhermack) were used to make the Polyvinyl Siloxane (PVS) samples for tensile testing validation experiments. The base and the catalyst were manually mixed 1:1 volume ratio for approximately 1 minute and cured at room temperature in 3D-printed dumbbell-shaped moulds and glass capillary tubes (d_in_ = 0.58 mm) for testing on the Instron 5544 (Instron) machine and on our device, respectively. In both cases, it was ensured that the length-to-width ratio of each specimen was greater than 4:1 to minimise any shear effects during tensile testing. To conduct mechanical testing in similar conditions on our device and on the Instron machine, a specific 3D printed base was designed allowing unconstrained motion of the ball bearing glued to the test specimen at one end and housing of a standard flat-surface clamp to secure the test specimen at the other end.

Prior to testing, a baseline current was applied to prevent the specimen from hanging and to align the ball with the CCD sensor. The current was thereafter controlled by means of the embedded software, and increased to ensure a minimum 10% elongation, necessary to capture the low-strain elastic behaviour of the Elite Double specimens. Strain rates of 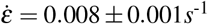 and 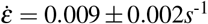 were used for the Elite Double 8 and 22 specimens, respectively. Three specimens per PVS formulation were tested three times apiece. The tensile force *F* applied on the sample could hence be calculated by means of the Equation 1. The engineering stress *σ* and the engineering strain *ε* were computed using 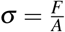 and 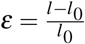, where *A* is the cross-sectional area of the sample, and *l* and *l*_0_ are the final and initial gauge lengths respectively. Testing of the three dumbbell-shaped specimens was carried out on the Instron machine using a 50.0 N load cell. Each sample was clamped at the two ends and stretched to a minimum of 10% strain at a comparable elongation rate 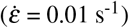 to that adopted during testing on our device. FEM (Abaqus) was leveraged to simulate the stress-strain behaviour of PVS samples attached at one end. Samples were modelled as cylinders of diameter d = 0.58 mm and of length 1_0_= 0.3 mm, composed of a perfectly elastic material with a Young’s modulus matching the literature value given in Table 1. Simulations were carried out for samples subjected to three different twist angles (*ϕ* = 0°, 22.5°, 45°), prior to applying uniaxial tension to 10% strain. Eight-node brick elements with reduced integration (C3D8R) were used for meshing. The von Mises stress σ_v_ was calculated at the mid-plane section of the object.

### Biological tissue samples preparation, testing experiments and modelling

All animal experiments were approved by the local ethical review committee. Work was approved by the University of Cambridge and conducted according to Home Office project licence PP0822646 at the Wellcome – CRUK Gurdon Institute, Cambridge University. A total of 12 wild-type C57BL/6J mice (Charles River; strain code 632) of approximately 12 weeks of age were used for this study. All animals were housed between 19 and 23°C, 45 and 65% humidity and a 12h day/12h night light cycle. All experiments comprised male and female mice, with no sex-specific differences.

After dissection from the body of the animal, the oesophagi were cut open, and the muscle layer was mechanically separated from the stroma and epithelium under a dissection microscope using watchmaker forceps as needed. Separation of the epithelial and stromal layers required a 2h incubation in PBS/100 mM EDTA at 37°C on an orbital shaker followed by mechanical separation under a dissection microscope using watchmaker forceps. All tissues were subsequently stored in PBS on ice (4°C), and tested within 6h on our device at room temperature (21 °C). Rectangular specimens were obtained via longitudinal cuts of the middle oesophagus (10 = 8.0 mm), using a 3D printed PLA cutting tool that was designed to the desired geometry and with slots to house razor blades at a fixed distance^43^. Final sample dimensions were measured under a dissecting microscope fitted with a calibrated eyepiece reticle. Absolute and relative thickness of the different tissue layers of the oesophagus wall were evaluated using oesophagus transversal tissue cryosections (7.0-10.0 *μ*m) stained with haematoxylin and eosin (H&E) (N=4 animals with 6 sections per animal) and imaged using an EVOS XL Core Imaging System.

A 3D printed PLA mounting support was used to guarantee the alignment of the wet tissue specimen during gluing with approximately 1 μL of cyanoacrylate (Loctite) to the chamber lids at one end and to the ball-to-specimen adaptor at the other end. Prior to testing, a baseline current was applied to align the ball and the specimen with the CCD image sensor. The current was controlled by the embedded software, and was progressively increased in a ramp-like fashion and strain rate was kept to a minimum, 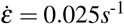, throughout test. At least three different test specimens were tested five times each for each condition. The tensile force *F* applied was computed per the Equation 1, and the engineering stress σ and engineering strain ε were computed as previously discussed.

A simple mechanical model, traditionally adopted to describe the behaviour of composite materials, was leveraged to evaluate the contribution of each individual tissue layer to the organ-level mechanical response as fully described in Supplementary Information.

## Data availability

Data associated with this study will be made openly accessible on public depositories at the time of publication.

## Code availability

The codes used for this study are available from the corresponding authors upon request and will be made openly accessible on public depositories at the time of publication.

## Acknowledgements

We are grateful to Benjamin D. Simons and his lab for insightful discussion and feedback on this work. This work was supported by the Engineering and Physical Sciences Research Council (EP/P021654/1). A.H. gratefully acknowledges the support of the University of Cambridge through a Herchel Smith Postdoctoral Research Fellowship and the support of Darwin College Cambridge through a Research Fellowship. He also acknowledges the support of the core funding to the Wellcome / CRUK Gurdon Institute (092096 and C6946/A14492) and of the Wellcome Grant (219478/Z/19/Z) to Benjamin D. Simons.

## Author contributions statement

L.R, A.H, L.C and T.S conceived the device and experiments. L.R and A.H conducted the experiments and analysed the data. A.H and T.S supervised the project. L.R, A.H and T.S wrote the manuscript. All authors reviewed the manuscript.

## Additional information

The authors declare no competing interests.

## Supplementary Figures

**Figure Sup.1.**
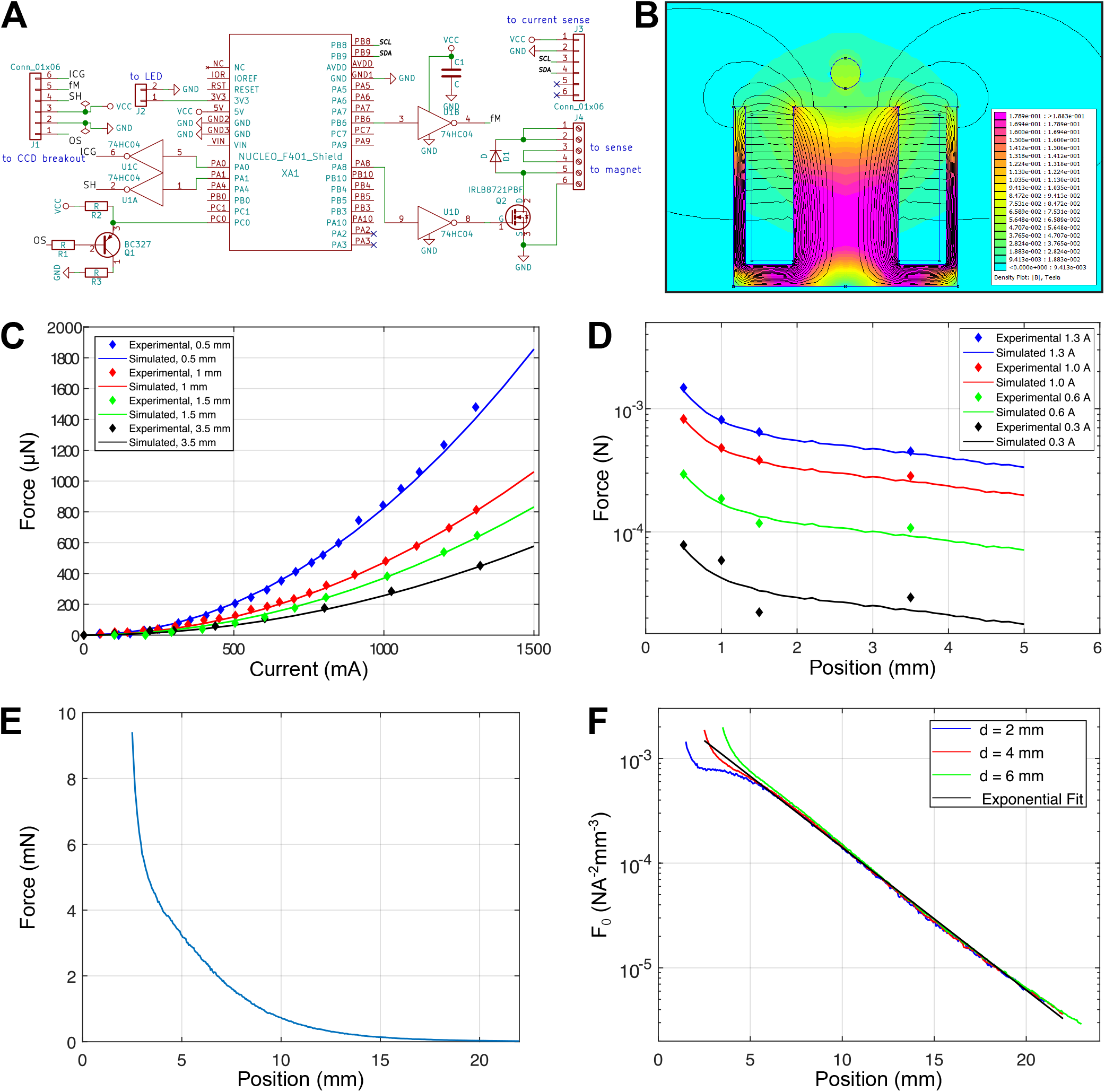
Characterisation of the magnetic force actuator. **A**: Schematic of the main printed circuit board attached to the headers of the microprocessor board, the magnet driver circuit and the breakout PCB attached to the CCD sensor. **B**: FEM simulation output. 2D heat map and contour lines of the magnetic field generated by the Eclipse M52173 electromagnet in presence of a 4.0 mm diameter bead made of 1010 grade mild steel. **C**: Simulated and experimentally measured values of magnetic force acting on the bead against current for a range of positions. **D**: Simulated and experimentally measured values of magnetic force acting on the bead against positions for a range of currents. **E**: Simulated magnetic force acting on a 4.0 mm bead at *I* = 280.0 mA. **F**: Force coefficient, F_0_ for a range of *d* and *I* values showing its exponential scaling behaviour.

**Figure Sup.2.**
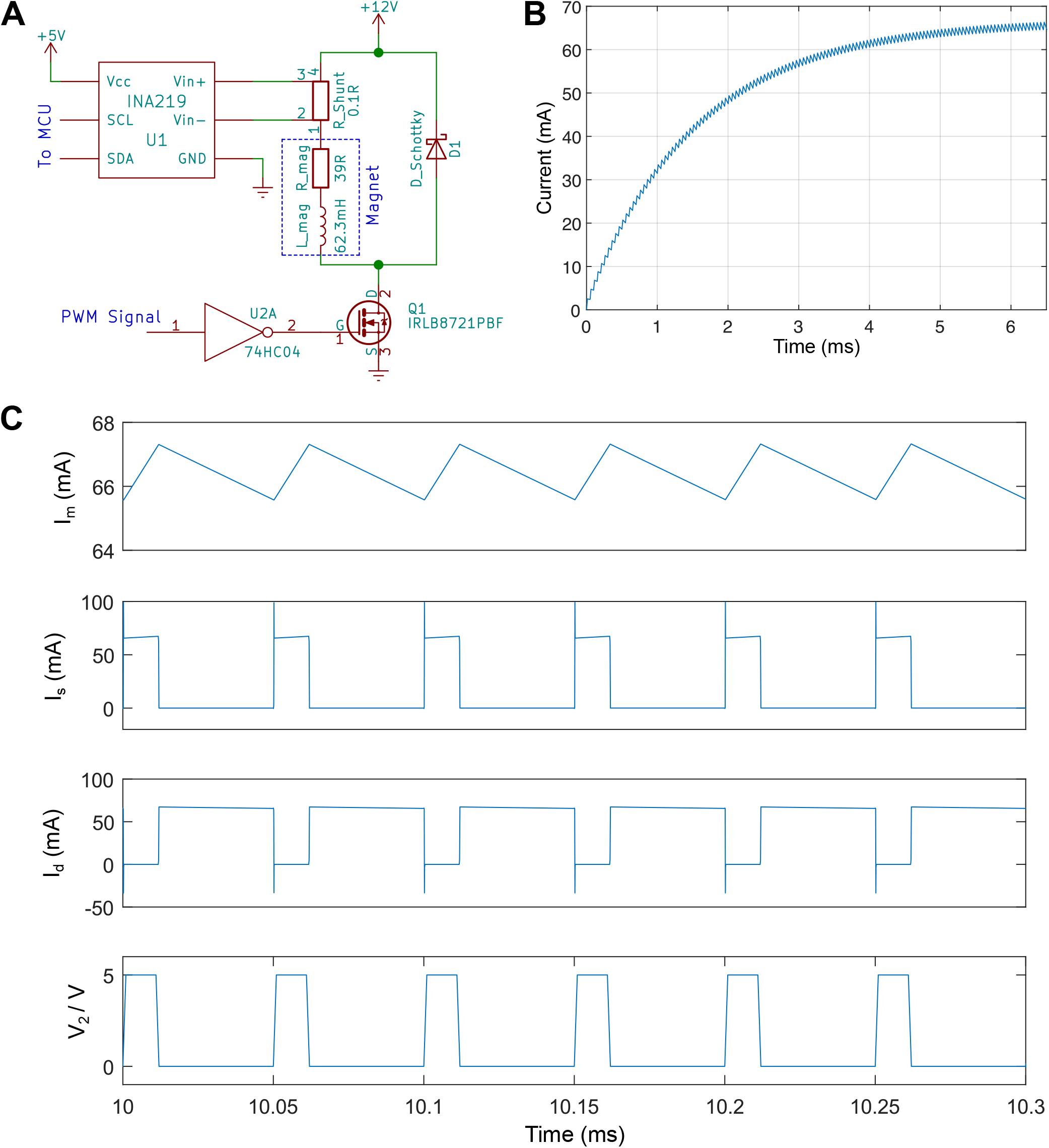
Design and characterisation of the electromagnet driver circuit. **A**: Schematic of the electromagnet driver circuit. **B**: Current transient response for a step change in duty cycle. **C**: Simulated current waveforms for the electromagnet, diode and power supply (*I_m_*,*I_d_*,*I_s_*) along with the driving 20% duty cycle, 20 kHz PWM signal. *I_m_* is the sum of *I_s_* and *I_d_*.

**Figure Sup.3.**
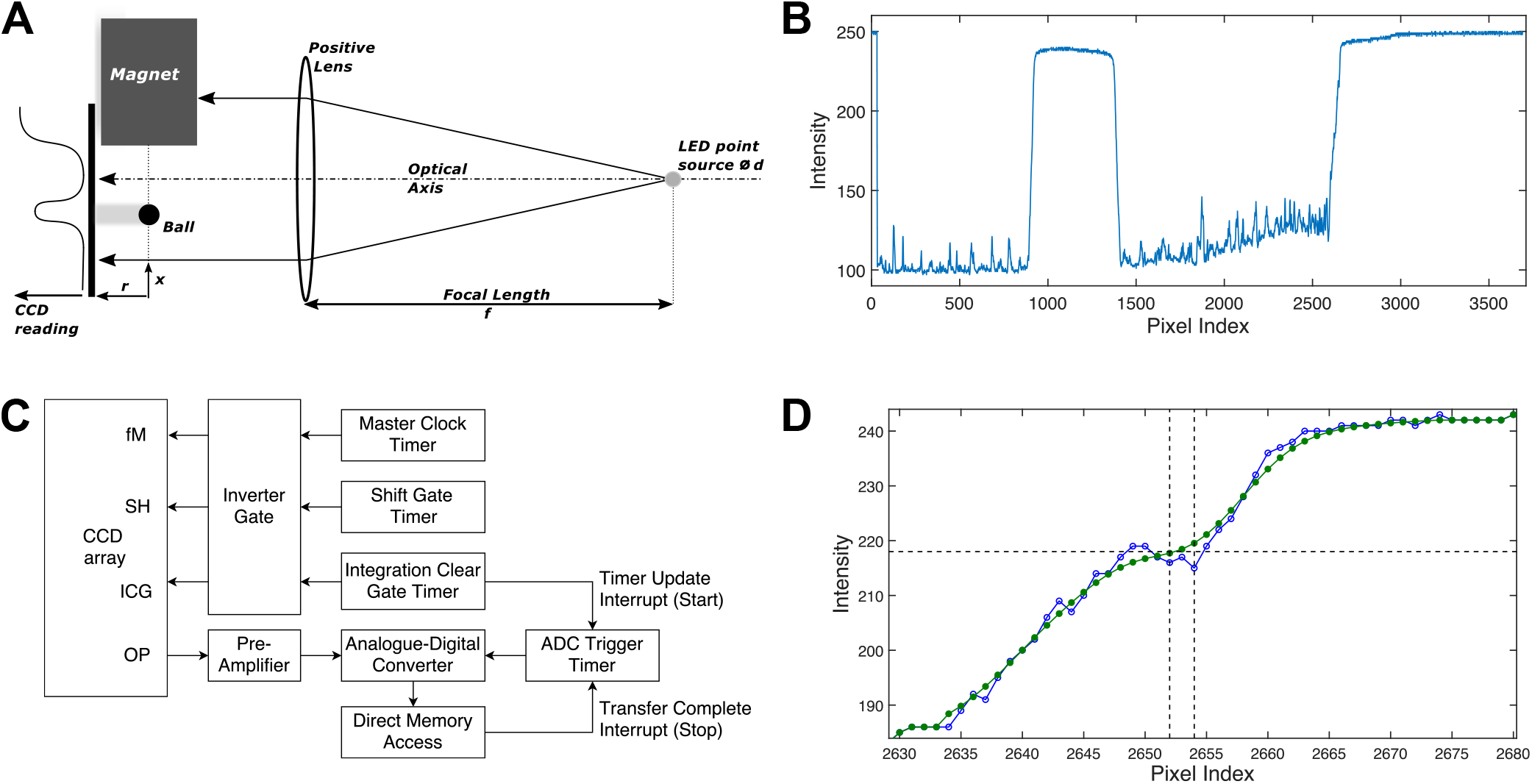
Design and characterisation of the CCD based optical sensor and associated embedded system. **A**: Schematic of the optical setup version using a lens and a LED point light source. **B**: Typical CCD output with high output corresponding to dark pixels associated with the cast shadow. On the right, one can observe the shadow of the edge of the electromagnet and, in the region between 900 and 1400 pixels, the shadow of the steel bead and buoyancy chamber. **C**: Schematic of the CCD driver architecture. **D**: Raw (blue) and filtered (green) CCD output signals at the edge of the magnet shadow. The threshold is shown as a horizontal line, and crossing points for the raw and smoothed signals are the right and left vertical lines respectively.

**Figure Sup.4.**
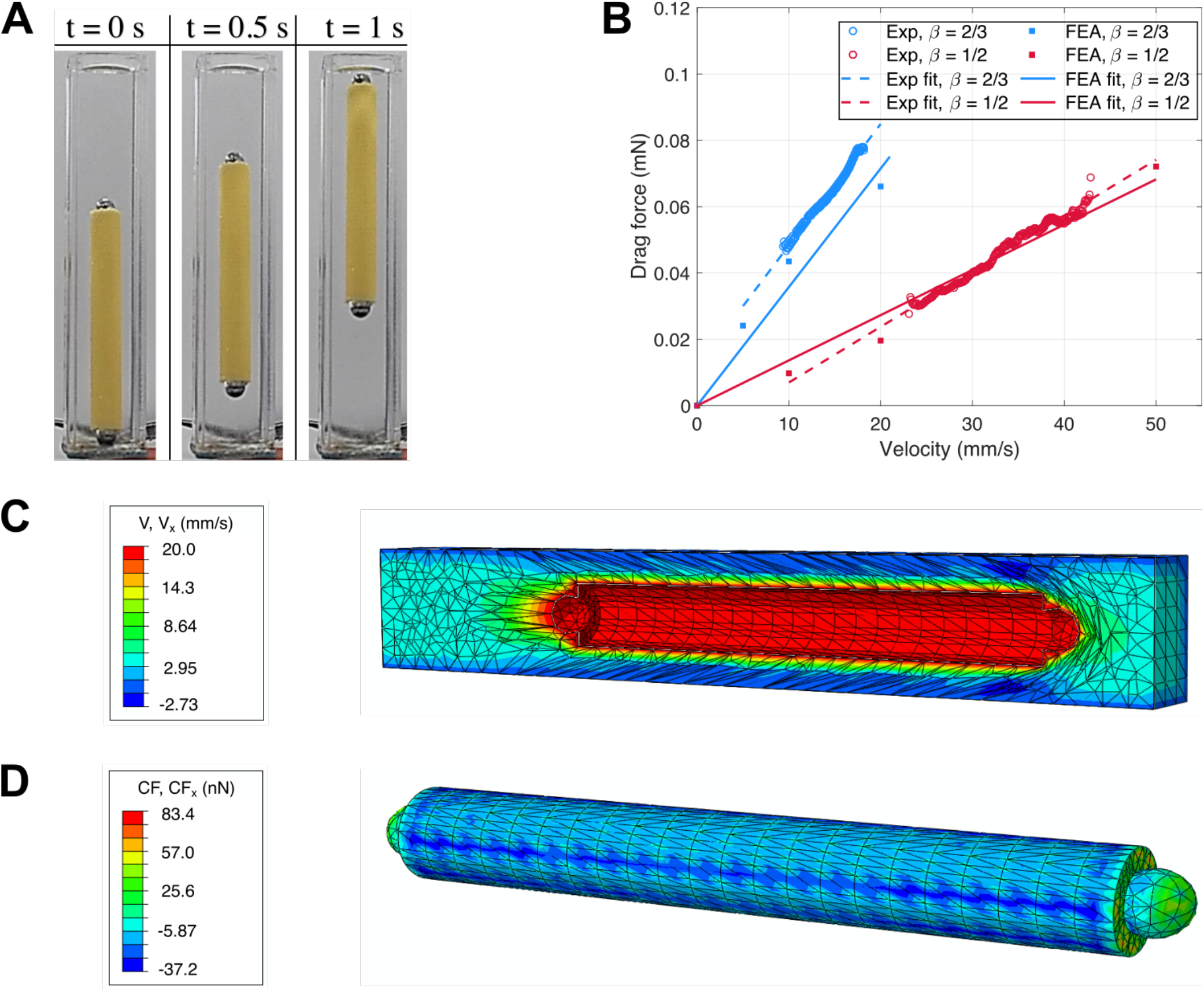
Characterisation of the drag force acting on the buoyancy cell. **A**: Experimental setup illustrating the position of the cell at *t* = 0,0.5 and 1s for 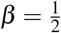. **B**: Force-velocity experimental and numerical results for 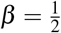 and 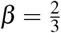. **C**: FEM results for the velocity profile for 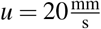 and 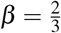. **D**: FEM results for the concentrated force on the buoyancy cell for 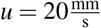 and 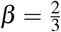.

**Figure Sup.5.**
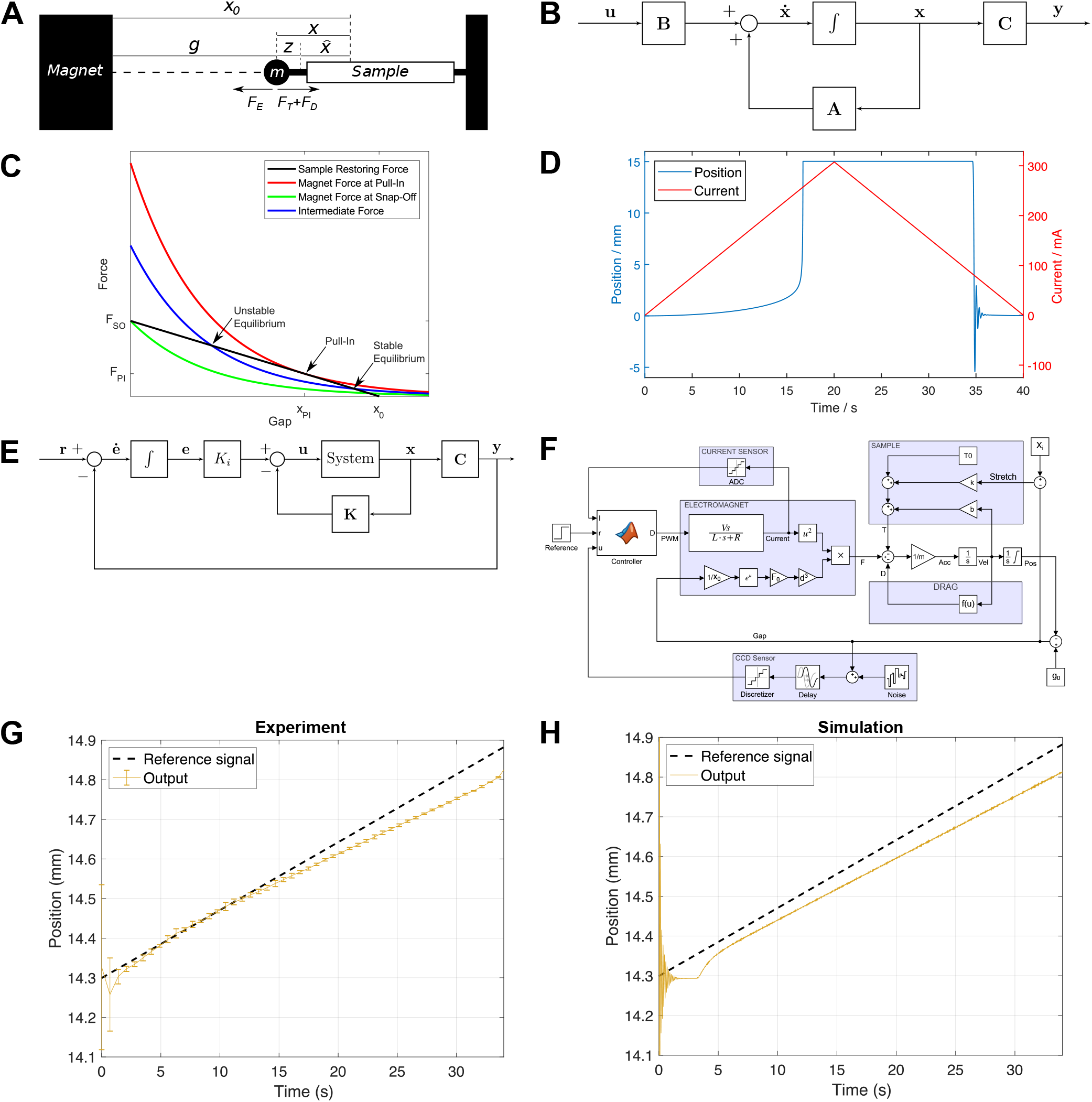
Mathematical model of the system and controller design. **A**: Schematic of the mechanical model of the system. **B**: Block diagram of the open-loop linearised system. **C**: Sample and magnet F(x) characteristics at critical and intermediate currents. **D**: Simulation of pull-in and snap-off events during a current sweep. **E**: Block diagram of a state-space controller configuration with integral action. **F**: Block diagram of the Simulink model of the system, highlighting the sample mechanics and drag force blocks. **G**: Experimental tracking of a reference input position signal. Error bars represent 1 SD. **H**: Simulated system behaviour for the same input.

**Figure Sup.6.**
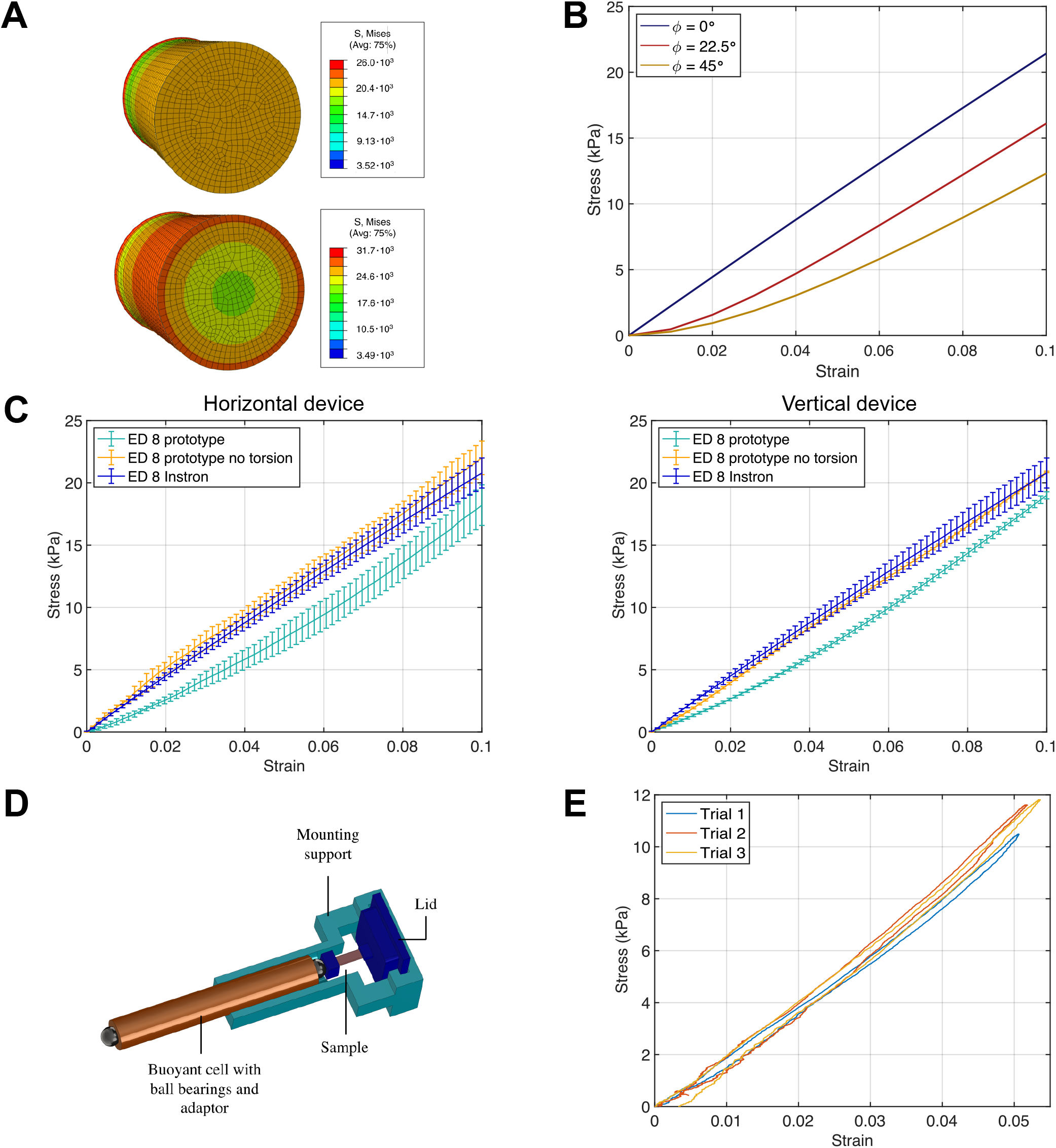
Device validation: effect of sample torsion, device configuration and hysteresis. **A**: FEM results showing the von Mises stress distribution at the mid-plane section for a cylindrical sample when no torsion is applied (top) and for a twisting angle of *ϕ* = 45°(bottom). **B**:Stress-strain curves computed numerically for three different twisting angles (*ϕ* = 0°, 22.5°, 45°). **C** Stress-strain curves for Elite Double 8 samples from the horizontal device configuration (air) with and without torsion effects (left). Stress-strain curves from the vertical device configuration under the same conditions. N=3 samples per condition. **D**: 3D Rendering of the mounting support allowing for an easy sample attachment and installation into the mounting chamber. **E**: Stress-strain curves for the device in horizontal configuration and using the mounting chamber filled with PBS and showing negligible hysteresis of one Elite Double 8 sample using a triangular reference input signal.

## Supplementary Information

### Magnetic force actuator driver circuit

A MOSFET-based switching circuit (Figure Sup. 2 A), was implemented to vary the input current of the electromagnet from a fixed DC voltage source. The input to the circuit is a pulse width modulated (PWM) square wave signal, which turns the MOSFET on for a fraction of the time, determined by the duty ratio D. To enable current to flow for the fraction of the cycle when the MOSFET is off, a Schottky diode is placed parallel with the electromagnet. Measurement of the current is achieved using a 0.1 **Ω** high-side shunt resistor and the Texas Instruments INA 219 current sensor, connected to a microprocessor over an I2C bus. The current sensor has a 12-bit analogue to digital converter, resulting in a current resolution of 0.1 mA, with a sampling period of 512 *μ*s.

Using LTspice circuit simulation software, we evaluated the response time and the current ripple of the driving circuit as shown in Figure Sup. 2 B and C. The response to a step change in D is consistent with that of a first-order inductor-resistor filter with a time constant *τ* = 1.6 ms. Under typical operating conditions of 12 V and 20 kHz modulation, the ripple is limited to 2.5 mA. It is important to minimise this disturbance due to the consequent unwanted oscillatory force applied to the sample and the effect on the accuracy with which the current may be instantaneously measured. Therefore, higher switching frequencies *f_PMW_* should be used, and when testing samples that require low forces, a smaller source voltage should be applied in preference to low duty cycles.

### Optical sensor embedded system

Accurate measurement of the extension of the sample necessitates accurate estimates of the distance between the rightmost edge of the bead shadow and the leftmost edge of the magnet shadow as shown in Figure Sup. 3 B. This can be done by using the pixel index at which the CCD signal drops below a certain threshold level and can be calculated with high precision. Figure Sup.3 D shows an example where the threshold level is crossed multiple times, leading to ambiguity in the choice of critical pixel index.

To increase the robustness of the measurement of the critical pixel index, we apply low-pass smoothing filter on the raw CCD signal. The filter is hard-coded as a Gaussian window function of length 15 pixels and only applied in the signal region in the vicinity of the threshold crossing through an iterative process. Iterating through the signal from the right (decreasing pixel indexes), the first crossing below the threshold value is identified (right vertical line in Sup.3 D). This gives the approximate location of the true crossing; in order to determine the precise value, the neighbouring ±20 pixels are filtered, using: 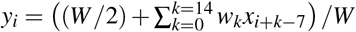, where *x_i_* and *y_i_* are respectively the raw and filtered signals, *w_k_* are integer filter coefficients,W is a normalising constant. The threshold crossing of the filtered 40-pixel signal can now be found to sub-pixel resolution using interpolation and is taken to be the true position of the magnet (the left vertical line in Figure Sup. 3 D). Continuing from the leftmost end of the filtered region, the same algorithm may be used to calculate the exact location of the right edge of the buoyancy cell and bead shadow, by identifying the point at which the filtered signal exceeds the threshold again.

The speed of this algorithm was optimised from an execution time of around 1.5 to 0.4 ms by adding a further coarse searching step, whereby every 50^*th*^ pixel is sampled to identify the 50-pixel block the transition lies in before applying the original algorithm to that block.

### Mathematical model of the device

The dynamics of the system, depicted in Figure Sup. 5 A, is governed by Newton’s Second Law, whereby the interaction between the electromagnet and the ferromagnetic bead is opposed by the tension in the sample and the fluid resistance, as described in Equations 3 and 4:

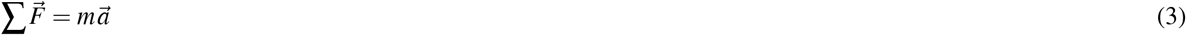

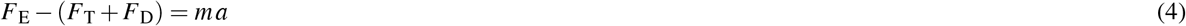

where *E*, *T*, and *D* refer to the electromagnetic, tensile, and drag forces respectively. This can be expressed in the standard state-space form:

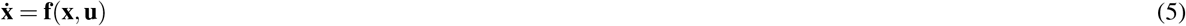

where **x** and **u** are the state and input vectors respectively, and **f** is the state transition function. Using Equation 4, and recalling from Equation 1 that the input to the system is the current *I* driving the electromagnet,**x** and **u** can be rewritten as:

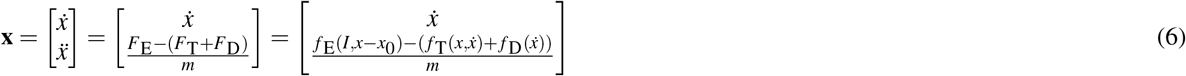

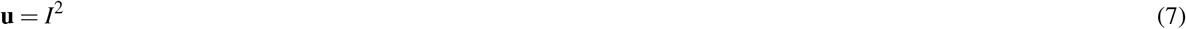

Rewriting this system in the standard linear form, as shown in the block diagram in Sup.5 B, yields:

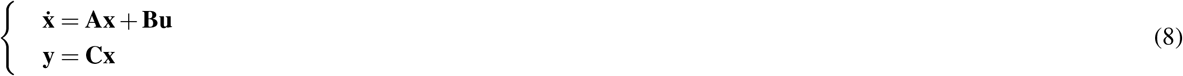

where the output of the system **y** is the position of the ferromagnetic bead measured by means of the optical sensor and it is related to the state vector **x** by the matrix **C**:

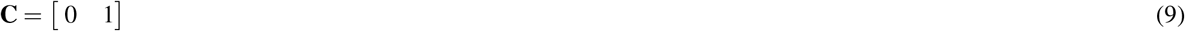

In order to compute matrices **A** and **B**, one must linearise the system in the vicinity of the equilibrium point 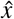:

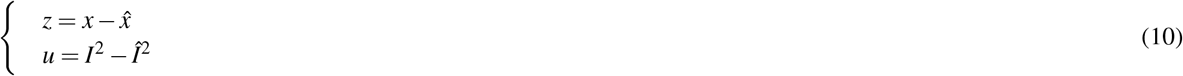

Computation of the first order Taylor expansion of **f** yields in the Jacobian matrix form:

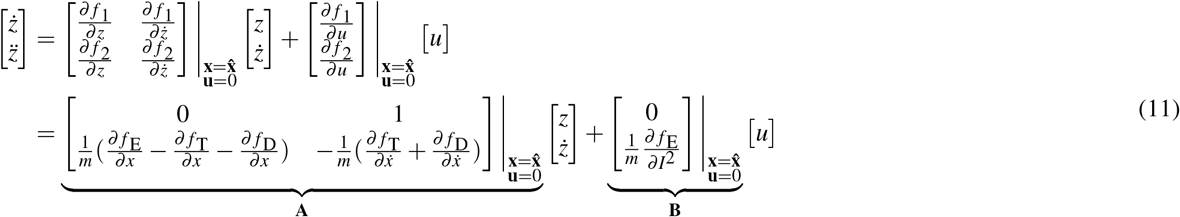

Coefficients of matrix **B** are only defined by the characteristics of the magnetic interaction between the bead and the electromagnet (Equation 1). However, coefficients of **A** also depends upon the drag force on the bead and buoyancy cell (Equation XX), and the tensile force generated by the test specimen. In order to derive this last force, we use a Kelvin-Voigt model consisting of a spring of stiffness, *k*, and a dashpot of viscosity, *b*, connected in parallel to model the viscoelastic behaviour of the tissue and add a constant force term *T*_0_ representing any component of the weight along the tensile direction:

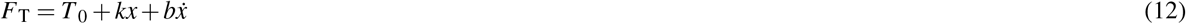

Rewriting the partial derivatives in Equation 11:

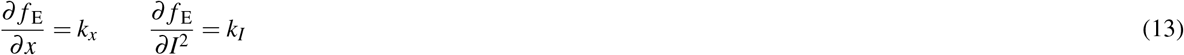

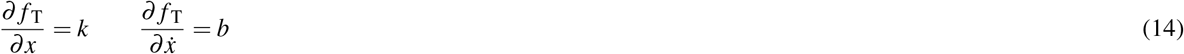

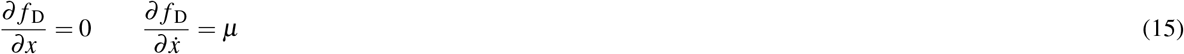

one can derive an analytical expression for matrices **A** and **B**:

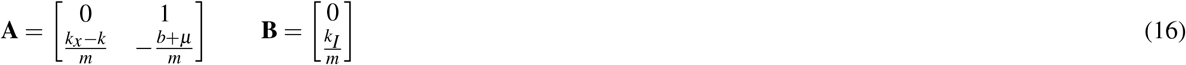

Here *k* and *b* are the parameters of the Kelvin-Voigt model, *k_x_* and *k_I_* are the so-called magnetic spring constants and *μ* is fluid viscosity constant. The stability of the open-loop system can now be assessed in deriving its poles from the characteristic equation (i.e. the eigenvalues of **A**):

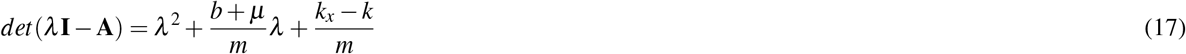

According to the Routh-Hurwitz criterion stability is achieved when the poles have negative real parts which is equivalent to the coefficients of the characteristic equation being positive. Thus, the system would be prone to instability for any *k_x_* > *k*, without the integration of a feedback loop.

Indeed, as shown in Figure Sup. 5 C the equilibrium points of the open-loop system occur when the magnet force balances the sample restoring force, *F_E_* = *F_T_*, neglecting the drag force in the first instance. Considering the point marked as stable equilibrium, where *k* > *k_x_*, a small deviation in ball position to the left leads to a greater increase in restoring force than in magnet force. The ball thus returns towards equilibrium; this is, by definition, stable. By contrast, a deviation to the left at the point marked unstable equilibrium leads to a greater increase in magnet force compared to the restoring force. The ball will therefore continue to accelerate towards the magnet. As the current is increased, a limiting case occurs when *F_T_* (*x*) becomes tangential to *F_E_*(*x*) with *k_x_* and *k* being equal. Beyond this marginally stable case, there is no equilibrium position at which the sample tension is equal to the magnet force, and the ball accelerates towards the magnet, an event termed pull-in. Stability cannot be achieved at positions greater than the pull-in location. Reducing the current below the pull-in value does not immediately release the ball; rather, the current must be reduced until the green line in Figure Sup. 5 C, at which point the ball accelerates towards the stable position near *x*_0_, an event termed snap-off. This behaviour is evidenced by the simulation in Figure Sup. 5 D, where hysteresis is clearly seen, with the current at pull-in being much greater than at snap-off (260 and 80 mA respectively).

### Controller design

The results of the analysis of the open-loop system clearly show the need for a controller. Without feedback, pull-in occurs at an extension as low as 30% of the initial gap, severely limiting the operating range of the instrument. To overcome this limitation, a proportional-integral-derivative (PID) controller was implemented. As shown in Figure Sup. 5 E, the controller takes as input the state vector **x** and the error between the reference **r** and the output **y**. Integration of the error with respect to time results in an internal integrator state **e** and the control signal **u**, which can be calculated according to:

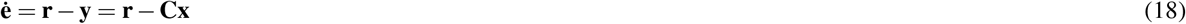

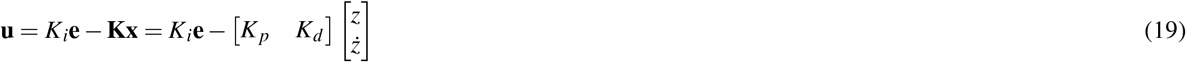

where *K_p_*, *K_i_*, and *K_d_* are, in order, the proportional, the integral, and the derivative gains. The proportional gain acts to modulate the current in opposition to the deviation of the ferromagnetic bead from the desired position; thus, if the bead moves closer to the electromagnet by, the square of the current is reduced by an amount *K _p_*, allowing the ball to return towards equilibrium. The role of the derivative term is to add damping to the response by opposing the velocity of the ball, while the integrator acts to prevent steady state error in the equilibrium position. Here, the state-space representation of the system becomes:

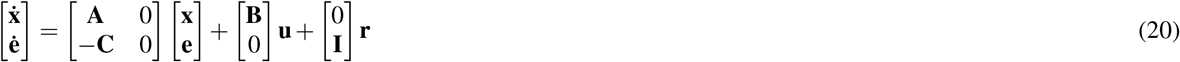

From combination of Equations 19 and 20, the closed-loop state transition matrix **A_*CL*_** can be defined as:

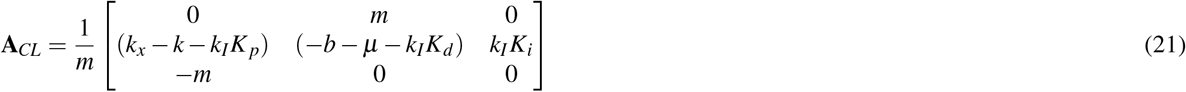

The poles of the closed-loop system are the eigenvalues of **A_*CL*_**, i.e. the roots of the characteristic polynomial below:

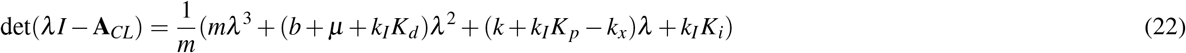

To compute the gains of the PID controller, this expression can be used in combination with the desired characteristic equation. This is obtained upon choice of the closed-loop poles *p*, as follows:

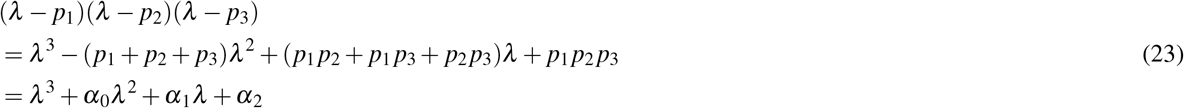

Therefore, the controller gains can be calculated by matching the coefficients of Equations 22 and 23. Assuming that *k*, *b*, and *μ* are zero in the first iteration yields:

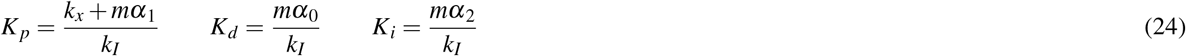

### Device simulation

As shown in Sup.5 F, the Simulink model consists of six main components, namely the *Electromagnet*, the *Sample*, and the *Drag* blocks, the current and image sensors, and the controller:

#### Electromagnet

The inputs to the first block are the distance between the metallic bead and the electromagnet as well as the duty cycle *D* of the pulse-width-modulated signal from the controller. This allows calculation of the current from:

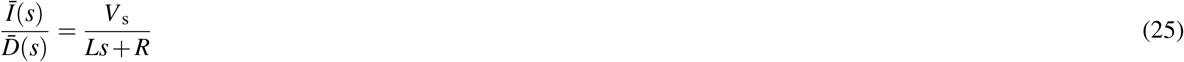

where *V*_s_ is the supplied voltage, and *L* and *R* correspond to the inductance and resistance of the circuit respectively. Thus, given a measure of the electric current, the electromagnetic force can be computed as per Equation 1.

#### Sample

Integration of the acceleration, which is calculated as per Equation 4, allows computation of the velocity and of the position of the bead. The elongation of the sample is therefore obtained, at each iteration, via comparison of the distance between the ball and the electromagnet with its initial value *x*_i_. The velocity and the elongation of the specimen are then multiplied by the viscosity and stiffness material constants respectively for calculation of the generated tension as per Equation 12.

#### Drag

The velocity of the sphere is utilised to compute the drag force as per 2. The combination of the electromagnetic force with the sample tension and drag resistance yields the net force acting on the sample (Equation 4).

#### Sensors

A current and image sensors are employed in the model, both providing inputs to the PID controller. The former measures the current from the *Electromagnet* as per Equation 25, whereas the latter records the distance between the sphere and the electromagnet and is modelled to add noise and delay to the system.

#### Controller

Alongside the inputs from the current and image sensors, the PID controller receives the reference position to be tracked, which, for the purposes of this work, was modelled as a ramp or a triangular signal. Its output is the duty cycle, which is in turn fed to the *Electromagnet* block for the following iteration to begin.

### Modelling soft biological tissues as composite materials

We propose a simple model allowing us to predict the mechanical behaviour of multi-layered biological tissues or organs given the response of each individual tissue layer. This model represents a generalisation of the *Rule of mixtures*, which is traditionally adopted to describe the behaviour of composite materials^44^. Our formulation is based on the analogous assumptions of ideal inter-layer bonding and uniaxial loading, whilst accounting for the non-zero residual stresses and strains of each individual layer. Nevertheless, it is valid exclusively in the regions of the stress-strain curve where the tissue exhibits an approximately linear elastic response.

Based on the tissue layers configuration exemplified in Figure 4 A, one can define a Representative Volume Element (RVE) of oesophagus wall as composed of epithelial (E), stromal (S), and muscle (M) layers. These are of variable length *l* and constant cross-sectional area *A*. When stress 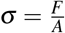 is applied to the RVE in the longitudinal direction, strain ε is generated as per 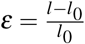. Generalising for *N* layers, from the principle of static equilibrium we obtain:

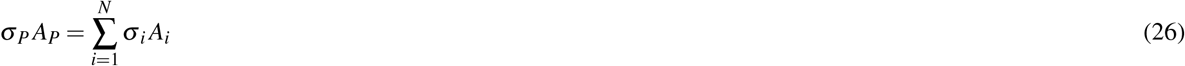

where the subscript *P* refers to the intact physiological state. The LHS and the RHS of the equation represent the intact and the zero-stress states respectively. For Equation 26 to hold, the physiological non-zero residual stress σ_*R*_ of each layer must be considered together with the stress due to stretching σ_*S*_, yielding:

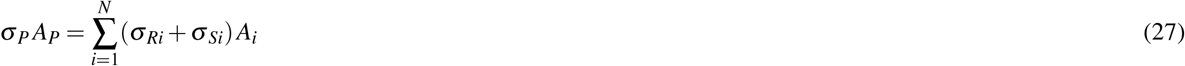

Dividing each side by *A_P_*, and for constant width, this can be rewritten as:

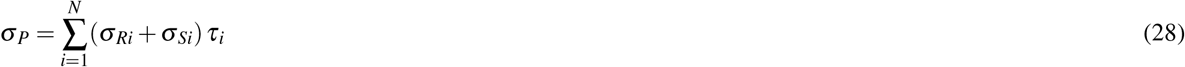

where *τ_i_* is the thickness fraction of each layer with respect to the intact RVE, such as 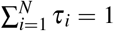. Thus, taking the first derivative of the stress with respect to strain, and considering non-zero residual strains on the RHS, we can derive an expression of the Young’s modulus in physiological state:

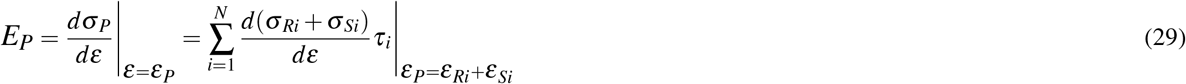

For constant values of σ_*R*_, this expression can be reduced to:

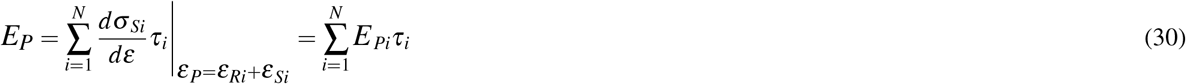

Thus, equation 30 relates the Young’s modulus, *E_P_*, of the intact oesophageal wall with that of each constitutive tissue layer in the zero-stress state, *E_Pi_*, determined at *ε_P_* = ε*_Ri_* + ε*_Si_*, where ε_*Ri*_ and ε_*Si*_ are respectively the residual strain and the *in vivo* stretch respectively. This formulation represents a weighted sum of each contribution, whereby, on the RHS, *E_Pi_* is translated by the residual strain of each layer, the *in vivo* stretch being the same for all tissue layers.

## References

1. Guimarães, C. F., Gasperini, L., Marques, A. P. & Reis, R. L. The stiffness of living tissues and its implications for tissue engineering. Nat. Rev. Mater. 5, 351–370, DOI: 10.1038/s41578-019-0169-1 (2020).

2. Hayward, M.-K., Muncie, J. M. & Weaver, V. M. Tissue mechanics in stem cell fate, development, and cancer. Dev. Cell 56, 1833–1847, DOI: 10.1016/j.devcel.2021.05.011 (2021).

3. Savin, T. et al. On the growth and form of the gut. Nature 476, 57–62, DOI: 10.1038/nature10277 (2011).

4. Hannezo, E. & Heisenberg, C.-P. Mechanochemical feedback loops in development and disease, DOI: 10.1016/j.cell.2019.05.052 (2019).

5. Hallou, A. & Brunet, T. On growth and force: mechanical forces in development. Development 147, DOI: 10.1242/dev.187302 (2020).

6. McGinn, J. et al. A biomechanical switch regulates the transition towards homeostasis in oesophageal epithelium. Nat. Cell Biol. 23, 511–525, DOI: 10.1038/s41556-021-00679-w (2021).

7. Vining, K. H. & Mooney, D. J. Mechanical forces direct stem cell behaviour in development and regeneration. Nat. Rev. Mol. Cell Biol. 18, 728–742, DOI: 10.1038/nrm.2017.108 (2017).

8. De Belly, H., Paluch, E. K. & Chalut, K. J. Interplay between mechanics and signalling in regulating cell fate. Nat. Rev. Mol. Cell Biol. 23, 465–480, DOI: 10.1038/s41580-022-00472-z (2022).

9. Fung, Y.-C. Biomechanics: Mechanical properties of living tissues. (Springer; New York, NY, Berlin, 1993), 2nd edn.

10. Gregersen, H. Biomechanics of the Gastrointestinal Tract (Springer London, 2003).

11. Phillip, J. M., Aifuwa, I., Walston, J. & WirtZ, D. The mechanobiology of aging. Annu. Rev. Biomed. Eng. 17, 113–141, DOI: 10.1146/annurev-bioeng-071114-040829 (2015).

12. Butcher, D. T., Alliston, T. & Weaver, V. M. A tense situation: forcing tumour progression. Nat. Rev. Cancer 9, 108–122, DOI: 10.1038/nrc2544 (2009).

13. Harn, H. I. et al. The tension biology of wound healing. Exp. Dermatol. 28, 464–471, DOI: 10.1111/exd.13460 (2017).

14. Cox, T. R. & Erler, J. T. Remodeling and homeostasis of the extracellular matrix: implications for fibrotic diseases and cancer. Dis. Model. & Mech. 4, 165–178, DOI: 10.1242/dmm.004077 (2011).

15. Gerhard A. Holzapfel, R. W. O. (ed.) Biomechanics of Soft Tissue in Cardiovascular Systems (Springer Vienna, 2003).

16. Walsh, M. et al. Uniaxial tensile testing approaches for characterisation of atherosclerotic plaques. J. Biomech. 47, 793–804, DOI: 10.1016/j.jbiomech.2014.01.017 (2014).

17. Akhtar, R., Sherratt, M. J., Cruickshank, J. K. & Derby, B. Characterizing the elastic properties of tissues. Mater. Today 14, 96–105, DOI: 10.1016/s1369-7021(11)70059-1 (2011).

18. Wu, P.-H. et al. A comparison of methods to assess cell mechanical properties. Nat. Methods 15, 491–498, DOI: 10.1038/s41592-018-0015-1 (2018).

19. McKee, C. T., Last, J. A., Russell, P. & Murphy, C. J. Indentation versus tensile measurements of Young’s modulus for soft biological tissues. Tissue Eng. Part B: Rev. 17, 155–164, DOI: 10.1089/ten.teb.2010.0520 (2011).

20. Griffin, M., Premakumar, Y., Seifalian, A., Butler, P. E. & Szarko, M. Biomechanical characterization of human soft tissues using indentation and tensile testing. J. Vis. Exp. DOI: 10.3791/54872 (2016).

21. Macrae, R. A., Miller, K. & Doyle, B. J. Methods in mechanical testing of arterial tissue: A review. Strain 52, 380–399, DOI: 10.1111/str.12183 (2016).

22. Harris, A. R. et al. Generating suspended cell monolayers for mechanobiological studies. Nat. Protoc. 8, 2516–2530, DOI: 10.1038/nprot.2013.151 (2013).

23. Savin, T., Shyer, A. E. & Mahadevan, L. A method for tensile tests of biological tissues at the mesoscale. J. Appl. Phys. 111, 74704 – 74704, DOI:https://doi.org/10.1063/1.3699176 (2012).

24. Landau, L. D. & Lifshitz, E. M. Fluid Mechanics (Pergamon, Oxford, England, 1987), 2nd edn.

25. Turki, S., Abbassi, H. & Nasrallah, S. Effect of the blockage ratio on the flow in a channel with a built-in square cylinder. Comput. Mech. 33, 22 – 29, DOI: 10.1007/s00466-003-0496-2 (2003).

26. Mercado-Perez, A. & Beyder, A. Gut feelings: mechanosensing in the gastrointestinal tract. Nat. Rev. Gastroenterol. & Hepatol. 19, 283–296, DOI: 10.1038/s41575-021-00561-y (2022).

27. Zhang, Y., Bailey, D., Yang, P., Kim, E. & Que, J. The development and stem cells of the esophagus. Development 148, DOI: 10.1242/dev.193839 (2021).

28. Colom, B. et al. Mutant clones in normal epithelium outcompete and eliminate emerging tumours. Nature 598, 510–514, DOI: 10.1038/s41586-021-03965-7 (2021).

29. Treuting, P. M. & Dintzis, S. M. (eds.) Comparative Anatomy and Histology (Academic Press, San Diego, 2012).

30. Goyal, R. K., Biancani, P., Phillips, A. & Spiro, H. M. Mechanical properties of the esophageal wall. J. Clin. Investig. 50, 1456–1465, DOI: 10.1172/jci106630 (1971).

31. Liao, D., Fan, Y., Zeng, Y. & Gregersen, H. Stress distribution in the layered wall of the rat oesophagus. Med. Eng. & Phys. 25, 731–738, DOI: 10.1016/s1350-4533(03)00122-x (2003).

32. Liao, D., Zhao, J., Fan, Y. & Gregersen, H. Two-layered quasi-3d finite element model of the oesophagus. Med. Eng. & Phys. 26, 535–543, DOI: 10.1016/j.medengphy.2004.04.009 (2004).

33. Lu, X. & Gregersen, H. Regional distribution of axial strain and circumferential residual strain in the layered rabbit oesophagus. J. Biomech. 34, 225–233, DOI:https://doi.org/10.1016/s0021-9290(00)00176-7 (2001).

34. Stavropoulou, E. A., Dafalias, Y. F. & Sokolis, D. P. Biomechanical and histological characteristics of passive esophagus: Experimental investigation and comparative constitutive modeling. J. Biomech. 42, 2654–2663, DOI: 10.1016/j.jbiomech.2009.08.018 (2009).

35. Gregersen, H., Lee, T. C., Chien, S., Skalak, R. & Fung, Y. C. Strain distribution in the layered wall of the esophagus. J. Biomech. Eng. 121, 442–448, DOI: 10.1115/1.2835072 (1999).

36. Yang, W., Fung, T. C., Chian, K. S. & Chong, C. K. 3d mechanical properties of the layered esophagus: Experiment and constitutive model. J. Biomech. Eng. 128, 899–908, DOI: 10.1115/1.2354206 (2006).

37. Zhao, J., Chen, X., Yang, J., Liao, D. & Gregersen, H. Opening angle and residual strain in a three-layered model of pig oesophagus. J. Biomech. 40, 3187–3192, DOI: 10.1016/j.jbiomech.2007.04.002 (2007).

38. Stavropoulou, E. A., Dafalias, Y. F. & Sokolis, D. P. Biomechanical behavior and histological organization of the threelayered passive esophagus as a function of topography. Proc. Inst. Mech. Eng. Part H: J. Eng. Medicine 226, 477–490, DOI: 10.1177/0954411912444073 (2012).

39. Holzapfel, G. A. Biomechanics of soft tissue. In Lemaitre, J. (ed.) Handbook of materials behavior models, 1057 – 1071, DOI:https://doi.org/10.1016/B978-012443341-0/50107-1 (Academic Press, Burlington, 2001).

40. Bancelin, S. et al. Ex vivo multiscale quantitation of skin biomechanics in wild-type and genetically-modified mice using multiphoton microscopy. Sci. Reports 5, DOI: 10.1038/srep17635 (2015).

41. Robinson, S. & Durand-Smet, P. Combining tensile testing and microscopy to address a diverse range of questions. J. Microsc. 278, 145–153, DOI: 10.1111/jmi.12863 (2020).

42. Jackson, J. D. Classical Electrodynamics (Wiley, 1998), 3rd edn.

43. Nelson, S. J., Creechley, J. J., Wale, M. E. & Lujan, T. J. Print-a-punch: A 3d printed device to cut dumbbell-shaped specimens from soft tissue for tensile testing. J. Biomech. 112, 110011, DOI: 10.1016/j.jbiomech.2020.110011 (2020).

44. Tsai, S. W. & Hahn, H. T. Introduction to composite materials (Technomic, 1980), 1st edn.

